# Identification of epigenetic modulators as determinants of nuclear size and shape

**DOI:** 10.1101/2022.05.28.493845

**Authors:** Andria Schibler, Predrag Jevtic, Gianluca Pegoraro, Daniel L. Levy, Tom Misteli

## Abstract

The shape and size of the human cell nucleus is highly variable amongst cell types and tissues. Changes in nuclear morphology are associated with disease, including cancer, as well as with premature and normal aging. Despite the very fundamental nature of nuclear morphology, the cellular factors that determine nuclear shape and size are not well understood. To identify regulators of nuclear architecture in a systematic and unbiased fashion, we performed a high-throughput imaging-based siRNA screen targeting 867 nuclear proteins including chromatin-associated proteins, epigenetic regulators, and nuclear envelope components. Using multiple morphometric parameters, we identified cell-type specific and general effectors of nuclear size and shape. Interestingly, most identified factors altered nuclear morphology without affecting the levels of lamin proteins, which are known prominent regulators of nuclear shape. In contrast, a major group of nuclear shape regulators were modifiers of repressive heterochromatin. Biochemical and molecular analysis uncovered a direct physical interaction of histone H3 with lamin A mediated via combinatorial histone modifications. Furthermore, disease-causing lamin A mutations that result in disruption of nuclear shape inhibited lamin A-histone H3 interactions. Finally, oncogenic histone H3.3 mutants defective for H3K27 methylation resulted in nuclear morphology abnormalities. Altogether, our results represent a systematic exploration of cellular factors involved in determining nuclear morphology and they identify the interaction of lamin A with histone H3 as an important contributor to nuclear morphology in human cells.

## INTRODUCTION

In physiological conditions, organelles have predictable morphology in a cell-type and tissue-specific fashion (Mukherjee et al., 2016). In contrast, abnormal and heterogenous organelle morphology is a prominent hallmark of many types of disease (Bexiga and Simpson, 2013; Galloway and Yoon, 2013). In particular, changes in nuclear size and shape are frequently associated with cancer and aging, and evaluation of aberrant nuclear morphology is routinely used in histology-based diagnostics (Cantwell and Dey, 2021; Mukherjee *et al*., 2016; Pathak et al., 2021; Zink et al., 2004). The comprehensive identification and characterization of cellular factors that determine and maintain nuclear morphology is important to elucidate the basic molecular mechanisms that determine overall nuclear architecture and to understand how abnormal nuclear morphology contributes to disease.

Nuclear morphology is highly plastic (Hoskins et al., 2021; Versaevel et al., 2012; Yoo et al., 2012). Changes to nuclear shape and size occur during developmental processes such as cellular division, differentiation, and migration (Skinner and Johnson, 2017). For example, in early *Drosophila melanogaster* development, embryonic nuclei appear spherical and small while at later stages they assume a more elongated shape with an increase in overall nuclear size. Mutant studies in *Drosophila* identified kugelkern (kuk), a lamin-like nuclear protein, as one factor required for these nuclear morphology changes (Brandt et al., 2006). In line with these nuclear shape changes during development, in adult tissues even within the same organism, different cell types often display distinct nuclear shapes and sizes (Mukherjee *et al*., 2016). The cellular factors that determine cell-type and tissue specific nuclear morphology are only partially understood.

One mechanism implicated in nuclear size control is transport through the nuclear pore complex (NPC) (Levy and Heald, 2012). The NPC acts as gateway between the cytoplasm and the nucleoplasm and both negative and positive regulators of nuclear transport have been shown to control the size of the nucleus (Jevtic et al., 2019; Levy and Heald, 2010; 2012). For example, *Tetrahymena thermophila* which is a unicellular eukaryote that maintains a macronucleus (MAC) and a micronucleus (MIC), expresses four Nup98 homologs which maintain MAC- and MIC-specific localization. Domain swapping experiments between MAC- and MIC-specific Nup98 proteins resulted in changes in nuclear size indicating that NPC composition can regulate nuclear size (Iwamoto et al., 2009). Similarly, the nuclear pore protein ELYS affects nuclear size in human epithelial cells by controlling NPC density and nucleocytoplasmic transport (Jevtic *et al*., 2019).

An obvious candidate determinant of nuclear morphology are nuclear lamins and the nuclear lamina (Deolal and Mishra, 2021; Pathak *et al*., 2021). The nuclear lamina is a proteinaceous network of multiple intermediate filament lamin proteins localized at the nuclear periphery between the nucleoplasm and the nuclear envelope. The human genome encodes three lamin genes: *LMNA*, *LMNB1*, and *LMNB2*, of which *LMNA* produces two protein isoforms, lamin A and lamin C (de Leeuw et al., 2018; Karoutas and Akhtar, 2021). Loss of or mutations in lamina proteins result in dysmorphic nuclei, increased DNA damage, and chromatin organization abnormalities and numerous human diseases, referred to as laminopathies (Marcelot et al., 2021; Shin and Worman, 2022; Wong and Stewart, 2020), which include striated muscle diseases, lipodystrophies, neurological syndromes, and premature aging disorders (Bonne et al., 1999; Kang et al., 2018; Karoutas and Akhtar, 2021; Mendez-Lopez and Worman, 2012). One dramatic laminopathy is Hutchinson-Gilford progeria syndrome (HGPS), an exceedingly rare aging disease which results in shortened lifespan, loss of subcutaneous fat, and cardiac abnormalities, among others (Gordon et al., 2014). HGPS is caused by a silent point mutation in *LMNA,* that leads to aberrant splicing and to the production of a mutant version of lamin A, referred to as progerin, which carries an internal 50 amino acid deletion at its C-terminus (Gordon *et al*., 2014). Progerin expression in HGPS cells acts in a dominant-negative fashion and results in misshapen nuclei, loss of heterochromatin, and increased endogenous DNA damage (Gordon *et al*., 2014). Furthermore, progerin has also been implicated in normal human aging (Scaffidi and Misteli, 2006).

In addition to the lamin proteins, chromatin has also been implicated in nuclear morphology. Early observations showed that in *Tetrahymena thermophila*, a specific nuclear histone linker protein is required for the reduced nuclear size in micronuclei (Allis et al., 1979; Shen et al., 1995). In addition, epigenetic readers and modifiers affect nuclear size. For example, overexpression of the histone H3 acetyltransferase BRD4 increases nuclear size in HeLa cells (Devaiah et al., 2016). In MCF-10A breast epithelial cells, a number of epigenetic and chromatin factors, including several core histones, have been shown to affect nuclear morphology (Tamashunas et al., 2020). Furthermore, recent observations suggest that the interaction of chromatin with lamins contributes to determining nuclear morphology (Karoutas et al., 2019; Stephens et al., 2019a). Single cell micromanipulation revealed two independent responses to mechanical forces, nuclei responded to small manipulations through chromatin and larger manipulations through lamin A/C (Stephens et al., 2017). Further studies of the role of chromatin in regulating nuclear morphology found chromatin to regulate nuclear dynamics and rigidity. In particular, manipulating the relative levels of euchromatin and heterochromatin altered nuclear architecture (Stephens *et al*., 2019a; Stephens et al., 2018). Furthermore, in cells with perturbed chromatin or lamins, increased heterochromatin suppressed nuclear blebbing and maintained nuclear rigidity (Stephens *et al*., 2019a; Stephens et al., 2019b). Close interplay between lamins and chromatin is also illustrated by the observation that loss of the lysine acetyltransferase MOF or its associated NSL-complex members KANSL2 or KANSL3 leads to altered mechanical properties of nuclei (Karoutas *et al*., 2019). While this effect appears to be due to reduced lamin acetylation, the observed changes in nuclear morphology are accompanied by alterations of the epigenetic chromatin landscape (Karoutas *et al*., 2019). These observations strongly suggest that nuclear size and shape are not determined by a single factor but rather are the result of an intricate interplay of architectural nuclear proteins with chromatin.

Nuclear morphology changes are routinely observed in pre-neoplastic and malignant cancer tissues. For example, tumor cells commonly display nuclear morphology abnormalities compared to nuclei in surrounding tissue (Chow et al., 2012). Anecdotal observations have identified multiple factors implicated in nuclear morphology and misregulation, and some of these factors have been linked to oncogenesis. For instance, mutations or alterations in expression of components of the LINC complex (Linker of Nucleoskeleton and Cytoskeleton) such as SYNE1 and SYNE2, which localize to the nuclear envelope and connect to the nuclear lamina, result in misshapen nuclei (Luke et al., 2008; Zhang et al., 2007). Alterations to SYNE1 and SYNE2 have been observed in colorectal (Yu et al., 2015), lung (Ahn et al., 2014), breast (Zuo et al., 2011), glioblastoma (Masica and Karchin, 2011), and ovarian cancer (Doherty et al., 2010). In addition, misexpression or mislocalization of nuclear lamins have been documented in both cancerous cells and tissues (Broers et al., 1993; Moss et al., 1999). Furthermore, lamin A/C is overexpressed in colorectal cancers where it correlates with poor prognosis and overexpressing GFP-lamin A in colorectal cancer cells increases cell motility (Willis et al., 2008). In contrast, reduced lamin A/C expression is documented in carcinomas of the esophagus, as well as breast, cervical, and ovarian cancers (Prokocimer et al., 2006).

Despite the fundamental nature of nuclear morphology and its link to disease, our knowledge of nuclear components and mechanisms that regulate size and shape is very limited (Cantwell and Dey, 2021; Mukherjee *et al*., 2016). Here we have used an imaging-based high-throughput RNAi screen to systematically identify factors that affect nuclear size and shape in multiple cell lines. We find known and novel determinants of cell-type specific and general nuclear morphology and uncover a novel mechanism of lamin-chromatin interactions mediated via histone H3 and its epigenetic modifications as a critical modulator of nuclear morphology.

## MATERIALS AND METHODS

### Human cell culture

Previously described karyotypically normal hTERT immortalized dermal fibroblasts cells (Scaffidi and Misteli, 2011) were maintained at 37°C with 5% CO_2_ in Minimum Essential Medium containing 15% Fetal Bovine Serum, 100U/mL Penicillin, 100 μg/mL Streptomycin, 2 mM L-Glutamine, and 1 mM Sodium Pyruvate. MCF10AT1k.cl2 cells (Barbara Ann Karmanos Cancer Institute) (Dawson et al., 1996; Heppner and Wolman, 1999) were cultured at 37°C with 5% CO_2_ in DMEM/F12 media supplemented with 1 mM CaCl_2_, 5% horse serum, 10 mM HEPES, 10 μg/mL insulin, 20 ng/mL EGF, 0.5 μg/mL hydrocortisone, and 0.1 μg/mL cholera toxin.

### High-throughput screen

For high-throughput siRNA screening, 1,200 cells were seeded into each well of a CellCarrier-384 Ultra microplate (PerkinElmer) using a Multidrop Combi Reagent Dispenser (ThermoFisher). Cells were reverse transfected with siRNAs targeting specific genes at a 20 nM concentration and were grown for 72 hours at 37°C. The siRNA knockdown screen used custom libraries (siRNA Silencer Select, ThermoFisher) targeting proteins that localize to the nuclear membrane (346 genes) or proteins involved in epigenetic and chromatin regulation (521 genes) (Table S1), each gene was targeted with 3 individual siRNAs placed in individual wells equaling 2601 siRNA experiments per screen. A non-targeting, scrambled siRNA (ThermoFisher, #4390847) was used as a negative control and the AllStars Hs Cell Death Control siRNA (Qiagen) was used as a control to score transfection efficiency and for assay optimization. An siRNA targeting lamin A/C was used as a positive biological control to score nuclear shape changes. After treatment, cells were fixed in 4% PFA in PBS for 20 min at room temperature, washed 3 times for 5 min in PBS, permeabilized with 0.5% Triton X-100 in PBS for 15 min, washed 3 times for 5 min in PBS, and blocked in PBS with 0.05% Tween 20 (PBST) and 5% BSA for 30 min. For detection and measurement of nuclear morphology, cells were immuno-stained with primary antibodies against lamin A/C (Santa Cruz, sc-376248, mouse, 1:1000) and lamin B1 (Santa Cruz, sc-6217, goat, 1:500) in PBST with 1% BSA for 4 hours at room temperature or overnight at 4°C. Cells were then washed 3 times for 5 min with PBST and incubated for one hour at room temperature with secondary antibodies diluted in 1% BSA in PBST containing DAPI (5 ng/μL), before washing 3 times for 5 min in PBST. The screen was performed in two biological replicates on different days.

Immuno-stained plates were imaged using an Opera QEHS (PerkinElmer) dual spinning disk high-throughput confocal microscope using a 40X water immersion lens (N.A. 0.9). The high-throughput microscope acquired images using 2 CCD cameras (1.3 Megapixels). The DAPI channel utilized the 405 nm laser for excitation and the 450/50 nm bandpass for acquisition, lamin B1 expression was imaged using the 488 nm laser for excitation and a 520/35 nm bandpass filter for acquisition, and lamin A/C expression was imaged using the 561 nm laser for excitation and the 600/40 nm bandpass filter for acquisition. DAPI, lamin B1, and lamin A/C were acquired at a single focal plane with pixel binning 2 (pixel size is equal to 323 nm). Thirty randomly selected fields were imaged per well and typically >250 cells per well were analyzed.

Quantification of nuclear size and shape were performed using a customized software pipeline. Images generated were analyzed using Columbus 2.6 software (PerkinElmer). An analysis pipeline was generated where nuclei were segmented using the DAPI staining image. Partial nuclei located at the edge of the image were excluded from subsequent steps of the analysis. Nuclear area, width, length, and circumference were measured as well as mean fluorescence intensity of lamin A/C and lamin B1. Single cell measurements were computed into mean per well values. RStudio software, R, and the cellHTS2 package (v 2.36.0) used the B-score method to normalize mean per well values on a per plate basis using the median of all the library wells on the plate. B-score values for all the samples in a single biological replicate were normalized to generate z-scores. The screen was performed twice on different days, and z-scores from each biological replicate were used to generate a final mean z-score. Hits were identified by z-scores +/-1.5 in at least 2 of the 3 unique siRNAs targeting each gene. Hits that fit these criteria but displayed a cell number with a z-score below −2 were also excluded to eliminate cell cycle or cell death abnormalities.

### Recombinant protein expression and purification

Fragments of human lamin A, C and progerin were cloned into the pGEX4T1 vector to generate N-terminally tagged GST fusion proteins. The plasmids used for recombinant protein production are listed in Table S2. Proteins were expressed in *Escherichia coli* Rosetta 2 (Novagen) in LB media. Protein expression was induced by the addition of 0.2 mM IPTG for 18 hours at 18°C. Cells were collected and suspended in lysis buffer containing 50 mM Tris pH 7.5, 150 mM NaCl, 0.05% NP-40, 1 mM PMSF, protease inhibitors, and 0.5 mg/mL lysozyme. Cells were incubated for 30 min on ice and lysed and sonicated for a duration of 20 seconds at 18% amplitude using a Branson Digital Sonifier 250 with a 102C converter set. Lysed cells were centrifuged at 21, 000 x g at 4°C for 15 min. The supernatant was removed and incubated with glutathione Sepharose 4B resin (GE). Beads were washed twice with lysis buffer and once with elution buffer (100 mM Tris-HCl, pH 8.0). Recombinant GST fusion proteins were eluted by resuspending the resin in elution buffer containing 15 mg/mL reduced L-glutathione (Sigma) and incubated at 4°C for 4 hours. Recombinant proteins were run on a 4-12% BisTris gel and analyzed using colloidal staining.

### Calf thymus histone binding assay

Calf thymus histone binding assays were performed by incubating 50 μg of calf thymus histones (Worthington) with 10 μg of purified GST fusion proteins in binding buffer containing 50 mM Tris pH 7.5, 1 M NaCl, and 1% NP-40 overnight at 4°C. To identify histone-lamin interactions, glutathione Sepharose 4B resin (GE) was added for 1 hour. Beads were washed 5 times in binding buffer, resuspended in 4x Laemmli buffer, run on a 4-12% BisTris gel, and transferred to a PDVF membrane. Membranes were used for western blot analysis using the antibodies at the specified concentrations (Table S3).

### Peptide array binding assay

MODified histone peptide arrays (Active Motif) composed of 384 unique histone peptides representing acetylation, citrullination, methylation, and phosphorylation posttranslational modifications were blocked with TBST (10 mM Tris/HCl pH 7.5, 0.05% Tween-20 and 150 mM NaCl) and 5% nonfat milk overnight at 4°C. The peptide array was washed twice with TBST, one time with interaction buffer (100 mM KCl, 20 mM HEPES pH 7.5, 1 mM EDTA, 0.1 mM DTT and 10% glycerol). The array was incubated with 10 nM purified GST-lamin A (pACS37) in interaction buffer at room temperature for 1 hour. The array was then washed three times in TBST, and incubated with anti-GST (GE, #27-4577-01, 1:5000) for 1 hour at room temperature in TBST with 1% non-fat dried milk. The array was further washed 3 times in TBST with 10 min for each wash and incubated with HRP conjugated secondary antibody (Santa Cruz) for 1 hour at room temperature. The membrane was submerged in ECL solution (Amersham), imaged, and intensity data was quantified using Array Analyzer Software (Active Motif).

### Expression of histone mutants in human cells

Lentiviruses containing wild-type histone H3.1 and H3.3 histone constructs were generated with the pLenti6.3/V5-TOPO TA Cloning kit (ThermoFisher). Mutations were added to wild-type H3.1-V5 and H3.3-V5 sequences using the Quikchange II XL site directed mutagenesis kit (Agilent). Lentiviruses were transfected into HEK293T cells in combination with viral packing and envelope plasmids pSPAX (Addgene 12260) and pMD2.G (Addgene 12259), respectively, and allowed to incubate for 48 hours before virus harvesting. Viral supernatant was collected from the HEK293T cells and placed onto hTERT immortalized fibroblast cells and incubated for 24 hours. Fibroblast cells were selected for mutant histone expression by the addition of 5 μg/mL of blasticidin for 10 days, 1200 cells were seeded onto 384 well plates and were grown for 72 hrs at 37°C. After treatment, cells were fixed in 4% PFA in PBS for 20 min at room temperature, washed 3 times for 5 min in PBS, permeabilized with 0.5% Triton-100x in PBS or 15 min, washed 3 times for 5 min in PBS, and blocked in PBS with 0.05% Tween 20 (PBST) and 5% BSA for 30 min. For detection and measurement of nuclear morphology, cells were immuno-stained with primary antibodies against lamin A/C (Santa Cruz, sc-376248, mouse, 1:1000) and lamin B1 (Santa Cruz, sc-6217, goat, 1:500) in PBST with 1% BSA for 4 hours at room temperature or overnight at 4°C. Cells were washed 3 times for 5 min with PBST and incubated for one hour at room temperature with secondary antibodies diluted in 1% BSA in PBST containing DAPI (5 ng/μL). Cells were washed 3 times for 5 min in PBST. Cells were imaged using the CV7000 high-throughput microscope (Yokogawa) with a 20X air objective (NA 0.75) and a sCMOS 2550×2160 pixel (5.5 Mpixel) camera. Images were binned 2X2 (pixel size is 650 nm). Images taken on the CV7000 microscope were analyzed using the Columbus 2.8.1 software (PerkinElmer). A Columbus image analysis pipeline segmented nuclei using the DAPI image, and then measured nuclear parameters were nuclear area, width, length, circumference, and the intensity levels of stained proteins.

### Statistical analysis

Scatterplots displaying the relationship between two measurements were assessed using Spearman’s coefficient analysis. Nuclear morphology features in cells expressing mutant histones were analyzed by using a two sample Kolmogorow-Smirnov test to compare the distribution of nuclear morphology scores at the single cell level.

## RESULTS

### An imaging-based screen to identify determinants of nuclear size and shape

We developed an imaging-based RNAi screen to identify and characterize novel cellular factors that regulate nuclear morphology, particularly nuclear shape and size (Fig. 1A). We focused on nuclear proteins that may affect nuclear morphology via interactions that take place either at the nuclear membrane or within the nucleus. To this end, we screened karyotypically normal hTERT immortalized dermal fibroblasts against two siRNA oligos libraries targeting 346 proteins that localize to the nuclear membrane and 521 proteins involved in epigenetic and chromatin regulation, respectively (see Materials and Methods; Table S1). For each gene targeted in the library, immortalized fibroblasts were reverse transfected in 384-well format with 3 unique siRNAs in 3 separate wells. 72 hours after siRNA reverse transfection, cells were fixed, permeabilized, and immuno-stained with antibodies against lamin A/C and lamin B1, and with DAPI to visualize DNA and to assess changes to nuclear shape (see Materials and Methods) (Fig. 1A). High-content imaging datasets were generated on a high-throughput spinning disk confocal microscope and nuclear morphology was quantified using an automated image analysis pipeline (see Materials and Methods) (Fig. 1A). To simultaneously assess multiple aspects of nuclear morphology, nuclear length, width, area, and roundness were measured simultaneously for all samples, treated as independent parameters, and used for hit identification (Fig. 1A). The screen was performed in biological duplicates and results were highly reproducible between replicates (Fig. S1). Non-targeting siRNAs were used as negative controls and a siRNA with a lethal phenotype was used as a positive technical control to measure transfection efficiency (see Materials and Methods). In addition, an siRNA targeting *LMNA* was used as a positive control based on the previously demonstrated role *LMNA* plays in maintaining nuclear shape (Lammerding et al., 2004) and as a control for fluorescence intensity levels of immuno-stained lamin A/C. Hits were identified using a per gene median z-score threshold of +/-1.5 (see Materials and Methods). To eliminate hits due to cell death or altered cell-cycle behavior, we excluded any hits with a cell number z-score of less than −2. As expected, among these hits were spindle assembly checkpoint components *MAD2L1*, *BUB1*, and cell cycle related kinases *AURKA* and *AURKB* (Fig. S2). Given the obvious mitotic defects caused by siRNA knock-down of these factors, rather than effects on the shape of interphase nuclei per se, they were deprioritized for downstream characterization.

**Figure 1.**
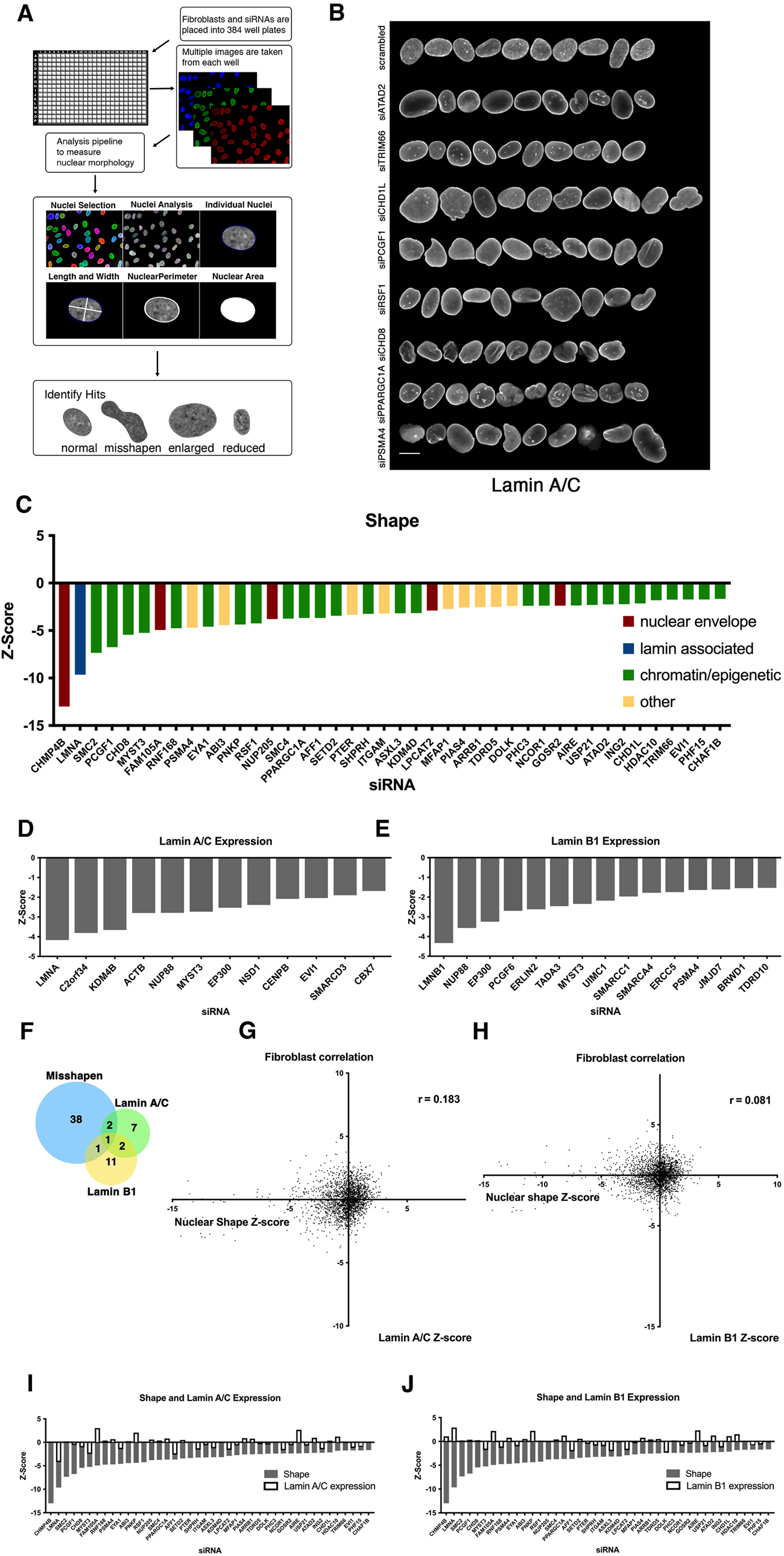
A high-throughput image-based screen for nuclear size and shape determinants. (A) Schematic overview of the nuclear morphology screen to identify genes required for proper nuclear shape and size. Cells were cultured in the presence of siRNAs targeting specific genes in 384-well plates, fixed, and stained with specific antibodies to visualize lamin A/C, lamin B1, or DAPI. Cells were imaged using high-throughput microscopy and an image analysis pipeline was developed to segment individual nuclei, measure nuclear morphology, and lamin A/C and lamin B1 intensity. Datasets were analyzed to identify misshapen, enlarged, and shrunken nuclei. Nuclear morphology changes were measured as z-scores. Typically, more than 250 nuclei were analyzed per sample. (B) Representation of normal nuclei and nuclear shape hits identified by high-throughput screening. Nuclear shape abnormalities are visualized by lamin A/C antibody staining. Scale bar: 10μm. (C) Nuclear shape hits were calculated by scoring circularity (circularity=4πArea/perimeter^2^). Z-scores were generated to compare hits across the screen. Nuclear shape hits were maintain a median z-score of −1.5. At least 250 nuclei were analyzed per sample. (D) Nuclear intensity of lamin A/C was assessed by calculating Z-scores of changes in lamin A/C expression on a per well basis. Lamin A/C hits were identified by median z-scores of −1.5 or less. (E) Nuclear intensity of lamin B1 was assessed by calculating median z-scores of changes in lamin B1 expression of −1.5 or less per well. (F) A comparison between lamin A/C and lamin B 1 expression hits and nuclear shape hits indicates lamin levels are not affected in most of nuclear shape hits. (G) Correlation between nuclear shape and lamin A/C expression the z-score of each parameter. Nuclear shape values relative to lamin A/C expression show little to no correlation. Spearman’s coefficient (r=0.183). (H) Scatterplot of nuclear shape values relative to lamin B1 expression show little to no correlation in lamin B1 protein expression levels and nuclear shape scores in fibroblasts. Spearman’s coeffiecient (r=0.081). (I) Nuclear shape hits in panel C with gray bars indicating the z-score for circularity. Lamin A/C z-score values are overlayed with white bars indicating lamin A expression changes. (J) Nuclear shape hits in panel C displaying z-score values. Lamin B1 z-score values are overlayed with white bars to indicating levels of lamin B1.

### Nuclear shape determinants

The phenotypic effect of siRNA knockdowns on nuclear morphology in the screen was quantified by Z-score analysis of the nucleus roundness parameter (roundness = 4Π Area / perimeter^2^, Fig. 1B, C; see Materials and Methods). Using this parameter, we identified 42 genes as positive hits in the primary screen, corresponding to a hit rate of 4.8 % (42/867, Fig. 1C). Nuclear shape regulators identified in the primary screen included genes whose products localize to the nuclear membrane or nuclear lamina such as *CHMP4B*, *LMNA*, or the nuclear pore component *NUP205*. Interestingly, 64% (27/42) of the effectors which altered nuclear shape were genes which are related to chromatin functions required for epigenetic modifications such as the polycomb component *PCGF1*, the histone acetyltransferase *MYST3*, the histone methyltransferase *SETD2*, or the ring finger protein *RNF168* (Fig. 1C).

Loss of lamins and lamin mutations have been linked to nuclear morphology changes in previous studies (De Sandre-Giovannoli et al., 2003; Eriksson et al., 2003; Lammerding *et al*., 2004). As expected, *LMNA* was identified as a prominent hit in our screen (Table S4A). To more broadly test how changes in lamin levels relate to alterations in nuclear shape, we mined our screening data for factors that lowered lamin A/C and lamin B1 levels (Fig. 1D-F). Twelve siRNA targets reduced lamin A/C level by a factor of at least 1.5 (Table S4B) and 15 targets reduced lamin B levels by 1.5-fold or more (Table S4C). Factors affecting both lamin A/C and lamin B1 levels include the nuclear pore protein *NUP88*, histone acetyltransferase *MYST3*, and the transcription co-factor *EP300*. To test whether the identified shape effectors exerted their effect on nuclear shape primarily via altering lamin levels, we assessed lamin A/C or lamin B1 in all shape effectors using quantitative imaging (Fig. 1I, J). Remarkably, of the 42 shape hits, only 4 affected lamin A/C and lamin B1 expression (Fig. 1F-J). This groups included the histone acetyltransferase *MYST3* which effects nuclear shape, lamin A/C, and lamin B1 levels, as well as the zinc-finger transcription factor *EVI1* and *LMNA* which affect both nuclear shape and lamin A/C levels, and the proteasome component *PSMA4* which affects nuclear shape and lamin B1 levels. However, in the majority of shape effectors (38/42), lamin A/C and lamin B1 levels were unaffected indicating that the majority of hits did not exert their effect on shape indirectly via altering lamin levels (Fig. 1F-1J). These results identify known modulators of nuclear shape, including factors which are accompanied by reduction of lamins, but more importantly, they point to a larger set of novel lamin-independent nuclear shape factors, including numerous epigenetic modifiers.

### Nuclear size determinants

In a complementary approach, we identified cellular factors that determined nuclear size. While nuclear size and shape are related features of nuclear morphology (Fig. 2A), we hypothesized that cellular factors exist that independently affect size or shape. Using the nuclear area as a proxy measurement for nuclear size (Fig. 2A), siRNA knock-down of 50 of the 867 genes affected nuclear size using a median z-score threshold of +/-1.5, corresponding to a hit rate of 5.7% (Fig. 2B, C). Amongst the size effectors, 36 increased nuclear size whereas 14 resulted in smaller nuclei (Fig. 2B, 2C). As observed for shape effectors, a large number of hits (52%; 26/50) were chromatin or epigenetic modifiers. Factors which increased nuclear size included genes encoding nuclear pore proteins such as *NUP205*, *NUP62*, *NUPL1*, and *NUP85* as well genes that encode components of histone modifiers such as *SUPT7L* and *PRMT2*. Factors that decreased nuclear size included the PhD-finger protein *PHF10*, the deacetylase *SIRT4*, the acetylation reader *BRD2*, and the histone methyltransferase *MLLT10* (Fig. 2B). Much like for nuclear shape, nuclear size determinates did not exert their effect via lamins, because of the 50 size determinants, none altered lamin levels (Fig. 2D).

**Figure 2.**
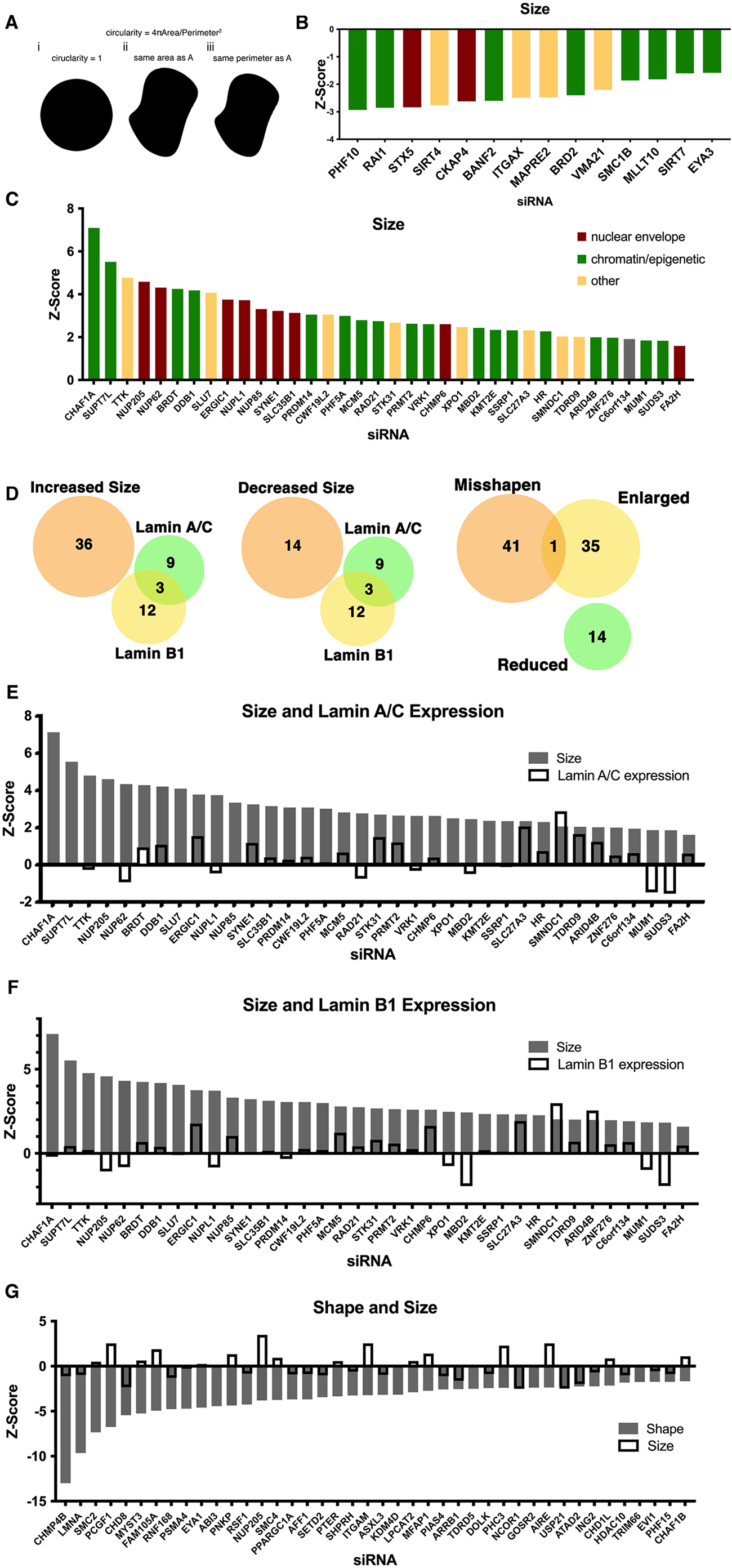
Identification of nuclear size determinants in human fibroblast cells. (A) A diagram of the relationship between circularity used to calculate nuclear shape and nuclear perimeter and area revealing the possibility that nuclear size might affect nuclear shape. Nuclear size hits were calculated by (circularity = 4πArea/perimeter^2^). Panel i shows a perfect circle with a circularity score of 1. Panel ii shows with the same area area as in panel A but an increased perimeter. Panel iii shows the same perimeter as panel A but a decreased area. If perimeter is a cellular constant, shape hits would correlate with a decrease in nuclear size. (B) Z-scores of hits displaying a decrease in nuclear size. Nuclear size hits maintain a median z-score of −1.5 or less. (C) Z-scores of hits displaying an increase in nuclear size. Nuclear size hits maintain a median z-score of 1.5 or greater. (D) Relationship of nuclear size hits and lamin A/C and lamin B1 hits. Nuclear shape hits show little overlap with nuclear size hits. (E) Nuclear size hits in panel C with gray bars indicating the z-score for hits displaying an increase in size. Lamin A/C z-score values overlayed with white bars indicating lamin A/C level. (F) Bar graph of nuclear size hits displaying z-score values. Lamin B1 z-score values are overlayed with white bars indicating levels of lamin B1. (G) Z-score values of nuclear shape hits in gray. Nuclear size z-score values are overlayed with white bars to show nuclear size varies among nuclear shape hits.

Given the relationship of nuclear size and shape, we cross-compared factors that affected both size and shape (Fig. 2D). Remarkably, we find almost no overlap between nuclear size determinants and nuclear shape effectors. Of the 42 shape effectors and 50 total size effectors, only one, the nuclear pore component *NUP205*, overlapped (Fig. 2D). These results suggest that nuclear shape and size are regulated by separate cellular pathways.

### Cell type specificity of nuclear morphology determinants

To understand nuclear morphology across multiple cell types and to ask whether nuclear size and shape determinants are cell-type specific, we repeated the screens in MCF10AT breast cancer cells (Fig. 3).

**Figure 3.**
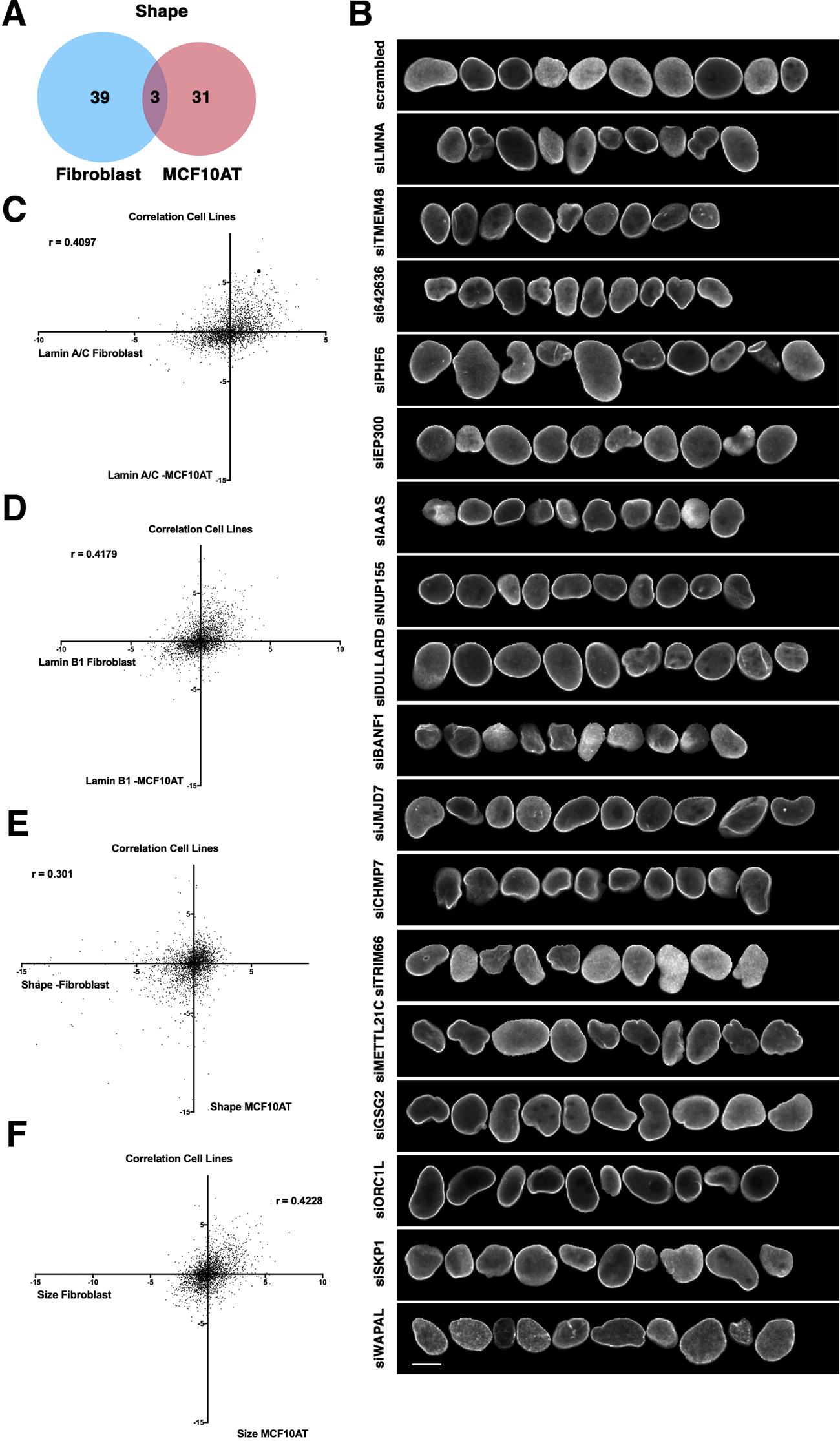
Cell-type specificity of effector hits. (A) Little overlap of hits for nuclear shape changes in immortalized human fibroblast cells compared to nuclear shape hits for the breast epithelial cell line MCF10AT. (B) Representative normal nuclei and nuclear shape hits of MCF10AT cells identified by high-throughput nuclear morphology screen and analysis. Nuclear shape abnormalities are visualized by lamin A/C antibody staining. Scale bar = 10 μm. (C) Lamin A/C z-score values in fibroblasts compared with lamin A/C z-score values in MCF10AT cells using the same siRNA target reveals little correlation between values. Spearman’s coefficient (r = .4097). (D) Lamin B1 z-score values in fibroblasts compared with lamin B1 z-score values in MCF10AT cells using the same siRNA target reveals little correlation between values. Spearman’s coefficient (r=.4179). (E) Nuclear shape z-scores of fibroblast and MCF10AT cell lines reveal a lack of overlap between the same siRNA targets. Spearman’s coefficient (r = .301). (F) Nuclear size z-score values in fibroblast cells compared with nuclear size z-score values in MCF10AT cells using the same siRNA target reveals little correlation between data points. Spearman’s coefficient (r = .4228).

Using the same measurement parameters, siRNA controls and image analysis pipeline as for fibroblasts, we identified 34 factors needed for proper nuclear shape in MCF10AT cells (Fig. 3A, B; Table S5). Interestingly, only three, *LMNA*, the transcription factors *EYA1* and *TRIM6,* overlapped with the shape determinants identified in fibroblasts (Fig. 3A, B). More generally, amongst all shape effectors in both screens, nuclear shape scores showed limited correlation (r = 0.301), suggesting nuclear shape is regulated via distinct pathways in fibroblast cells compared to breast epithelial cells (Fig. 3D).

Comparison of nuclear size effectors between fibroblasts and MCF10AT cells also showed little overlap (r = 0.4228) (Fig. 3F). Of the fibroblast and MCF10AT nuclear size hits only four overlapped (the chromatin assembly factor *CHAF1A*, DNA binding protein *DDB1*, transcription factor *PRDM14*, and transport protein *SLC27A3)* (Fig S5). While *SLC27A3* functions at the nuclear membrane, *CHAF1A, DDB1,* and *PRDM14* act through chromatin. Along with the cell-type specific effectors of nuclear size and shape, limited correlation between the effectors of lamin A/C and lamin B levels were found between cell types (Fig. 3C, D).

Taken together these results indicate that the mechanisms involved in the maintenance of nuclear size and shape are cell-type specific.

### Lamin A directly interacts with histone H3

Given our identification of numerous epigenetic modulators as determinants of nuclear size and shape combined with the lack of accompanying changes in lamins, we considered that chromatin-lamin interactions might mediate the observed size and shape effects. In particular, based on the identification in our primary screens of several post-translational modifiers of histone H3, including the histone H3K36-specific lysine methyltransferase *SETD2*, the histone H3K9 lysine demethylase *KDM4D*, and the histone deacetylase *HDAC10*, we hypothesized that histone H3-lamin interactions may contribute to maintaining nuclear size and shape.

To test this hypothesis, we first asked whether lamin A/C could bind directly to chromatin *in vitro*. Lamin A and lamin C consist of an N-terminal head domain followed by a long rod-like domain in the central region of the proteins and prior observations showed that the C-terminus of lamin A maintains an IgG-like fold domain and directly binds to DNA (Stierle et al., 2003). For that reason, and because many disease mutations affecting nuclear morphology localize to that region (Dittmer and Misteli, 2011; McKenna et al., 2013), we probed for a direct physical interaction of the C-terminal region of lamin A/C with chromatin (Fig. 4). We generated GST-fusions proteins of various fragments of lamin A or C, purified them, and incubated them with histones derived from calf thymus to test for direct binding to histones (Fig. 4A, B). We find that GST-lamin A containing the entire Ig-fold and C-terminus (aa 389-646) binds directly to histone H3, but not to the other core histones (Fig. 4B). GST-lamin A lacking the C-terminal 80aa (aa 389-566) or the C-terminal region of GST-lamin C (aa 389-572) did not bind to histone H3, suggesting that the unique portion of lamin A present in the C-terminus tail is required for the histone H3 interaction (Fig. 4B). However, while the lamin A tail was required for H3 binding, this region (aa 565-646) was not sufficient for binding (Fig. 4B), possibly due to its disordered nature (Qin et al., 2011). We conclude that lamin A can directly interact with core histone H3 via its C-terminal tail along with a portion of the homologous region present in both lamin A and lamin C.

**Figure 4.**
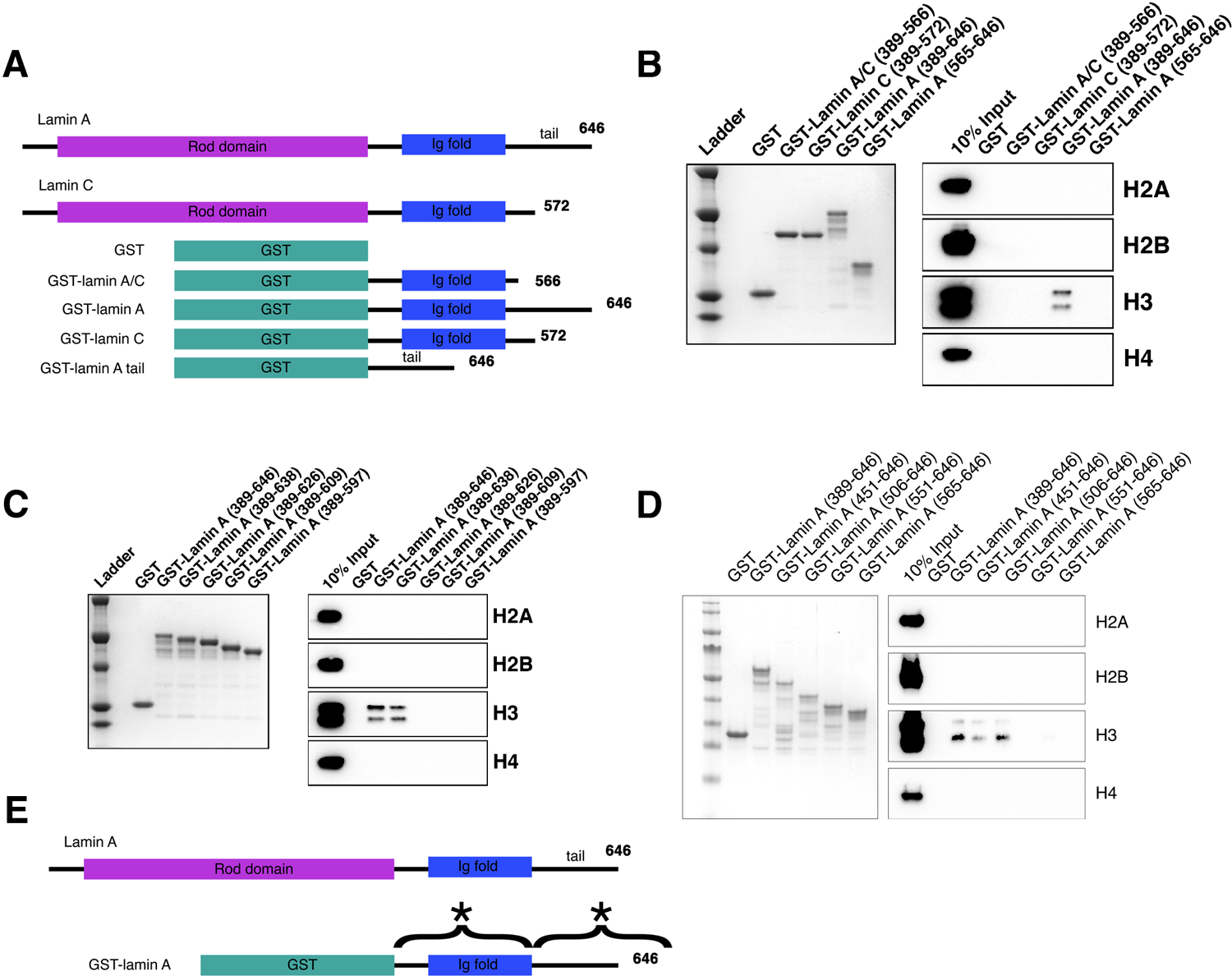
In vitro binding of lamin A to histone H3. (A) A diagram of constructs used in binding assays. (B) Colloidal staining of purified recombinant proteins and histone pull-down assays. GST-lamin A (389-646) directly binds to histone H3 but not histones H2A, H2B and H4. The portion of lamin A and C that is homologous (GST-lamin A/C (389-566), GST-lamin C (389-572) and the GST-lamin A truncation containing the C=terminal tail (GST-lamin A 565-646) did not bind histones. (C) Colloidal staining of purified recombinant proteins and histone pulldown assay. The C-terminal portion of lamin A is required for binding histone H3. Full length GST-lamin A (389-646) and the truncated GST-lamin A (389-638) bound histone H3 but not histones H2A, H2B, and H4. Further truncations to the lamin A tail (389-626), (389-609), and (389-597) did not interact with histones identifying the portion of the C-terminal tail essential for binding histone H3 as aa 638-646. (D) Colloidal staining of purified recombinant proteins and histone pulldown assay. The N-terminal portion of lamin A required for binding histone H3. Full length GST-lamin A (389-646) and the truncated GST-lamin A (451-646) and GST-lamin A (506-646) bound histone H3 but not histones H2A, H2B, and H4. Further truncations to the GST-lamin A (551-646) and GST-lamin A (565-646) did not interact with histones identifying the portion of the N-terminus essential for binding histone H3 as aa 506-550. (E) A schematic summary of the two regions required for lamin A-H3 interactions marked with (*).

To more precisely identify the region of the lamin A C-terminus required for the interaction with histone H3, multiple truncation mutants were generated and used in *in vitro* histone binding assays. GST-lamin A (aa 389-638) bound as well to histone H3 as the full-length lamin A tail (GST-lamin A, aa 389-646), while the aa 389-626 region did not (Fig. 4C). These experiments identify aa 627-638 as required for the lamin A-histone H3 interaction. Interestingly, this is the region deleted in the lamin A mutant isoform that causes the premature aging disorder HGPS, which is characterized by extensive nuclear shape aberrations including prominent nuclear lobulations and altered H3K9 and H3K27 methylation (Goldman et al., 2004; Shumaker et al., 2006). In line with a possible role of lamin A-histone H3 interactions in HGPS, GST-progerin (aa 506-664 D50) (Fig. S7A) did not bind H3 (Fig. S7B).

Binding of lamin A to naked DNA had previously been shown to occur in the context of lamin A dimers (Stierle *et al*., 2003). To ask whether dimerization is required for lamin A-H3 interactions we incubated GST-lamin A fusion proteins with the reducing agent DTT to inhibit dimerization. The addition of DTT increased lamin A-H3 interactions suggesting lamin A binds to histone H3 as a monomer and dimerization limits its binding (Fig. S7C). Furthermore, since both lamin A and lamin C can form dimers, bind to DNA, are co-expressed *in vivo*, and lamin C does not bind histone H3, we asked whether lamin C binding to lamin A can inhibit histone H3 binding via dimer formation. Co-incubation of the C-termini of GST-lamin A (aa 389-646) and GST-lamin C (aa 389-572) in a histone pulldown assay, reduced lamin A-histone H3 binding (Fig. 7B). The addition of DTT reversed inhibition, suggesting lamin C inhibits lamin A through dimerization at cysteine residues (Fig. S7D). These observations demonstrate direct interaction of lamin A with core histone H3.

### Interaction of lamin A with histone H3 is sensitive to epigenetic modifications

To further identify how lamin A binds to histone H3, we utilized a histone peptide binding array to probe the effect of histone tail modifications on lamin A-histone interactions (Fig. 5). The array consists of 384 unique histone peptides spotted onto a slide representing peptides of histones H2A, H2B, H3, and H4 with multiple common modifications including serine/threonine phosphorylation, lysine acetylation and methylation among others (see Materials and Methods). Binding assays of lamin A to the array confirmed interaction of lamin A with histone H3, and also revealed preferential binding to several histone modifications, including combinations of modifications (Fig. 5A; Table S6). In fact, the peptides that showed the strongest binding to lamin A contained a combination of modifications (Table S6). Specifically, lamin A bound preferentially to peptides which contained a methyl-methyl modification signature, including histone H3R8me2s/K9me2 (Fig 5B), H3K26me2s/K27me2 (Fig. 5C), and histone H4R19me2s/K20me1 (Fig. 5D). In line with this observation, GST-lamin A binding to peptides containing only a single modification, such as methylated arginine or methylated lysine alone, was reduced when compared to the three dually modified peptides (Fig. 5B-D; Table S6). For example, GST-laminA bound ∼ 5-fold more efficiently to histone H3R8me2s/K9me2 than histone H3K9me2. While most peptides with the strongest binding were dual methyl marks, we did find lamin A also bound to acetylated histone H3 and H4 peptides (Table S6). These include histone H4K12ac/K16ac, H3K27ac, and H3K4ac peptides although at reduced levels (Table S6). These data indicate methyl-methyl motifs represent the highest affinity target sites for lamin A-chromatin binding.

**Figure 5.**
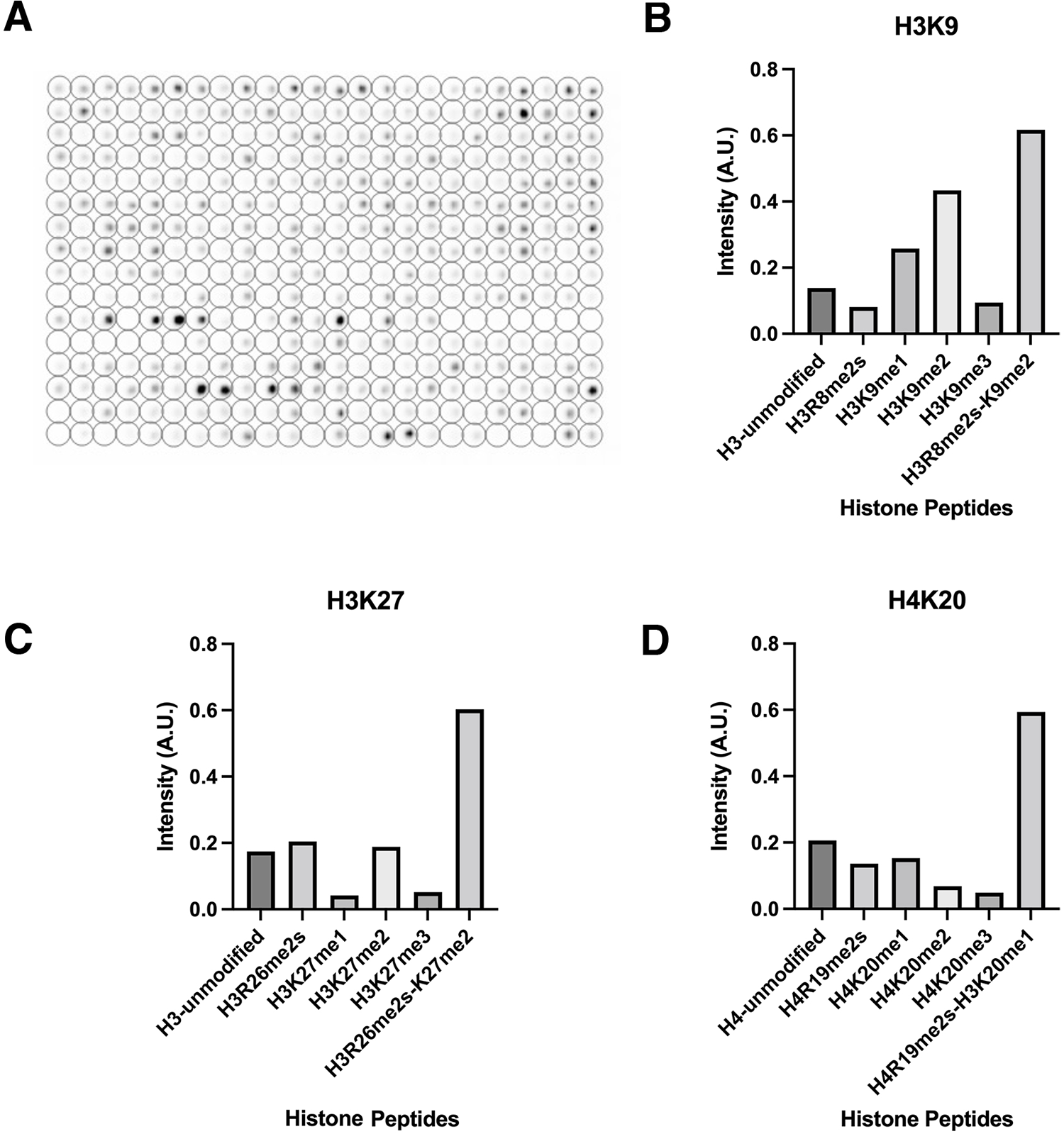
Specificity of lamin A binding histone modifications. (A) In vitro peptide binding array assay using GST-lamin A (506-646). Intensity of signal indicates binding. (B) Peptide binding assays for select histone H3K8/9 modifications. H3R8me2s/K9me2 maintained the most intense signal compared to single modifications alone. (C) Peptide binding assay for histone H3R26/K27 modifications. H3R26me2s/K27me2 maintained the most intense signal compared to single modifications alone. (D) Peptide binding assay for histone H4R19/K20 modifications. H4R19me2s/K20me1 maintained the most intense signal compared to single modifications alone. Values represent intensity.

### Histone H3.3 mutants result in nuclear shape and size abnormalities

The binding of lamin A to histone H3 is of interest since mutations in this histone variant have been implicated in disease. Mutations to histone H3.3 were first found in pediatric high grade glioma and later in chondrosarcomas and giant cell tumors of the bone (Weinberg et al., 2017). Recently, mutations to histone H3.3 have been associated with congenital disorders such as craniofacial and brain abnormalities, and developmental delay among others (Bryant et al., 2020). Interestingly, most of these diseases, including gliomas and chondrosarcomas, are characterized by changes in nuclear morphology (Nafe et al., 2003; Welkerling et al., 1996). We thus asked whether disease relevant H3 mutants are sufficient to induce nuclear morphology defects. Histone H3.1 variants carrying dominant mutations K9M, or K27M or H3.3 mutants at K9M, K27M, or K36M were stably expressed in hTERT immortalized fibroblast cells and nuclear morphology assessed. While histone H3.1 mutations had little or inconsistent effect on nuclear morphology, H3.3 mutants resulted in nuclear morphology changes and a decrease in nuclear size (Fig. 6; Fig. S8). Morphological changes displayed in H3.3 mutants were not due to cytotoxicity or proliferation defects, since no changes to overall cell number were detected. We conclude that histone H3.3 mutations involved in lamin A interactions contribute to dysmorphia of the human cell nucleus.

**Figure 6.**
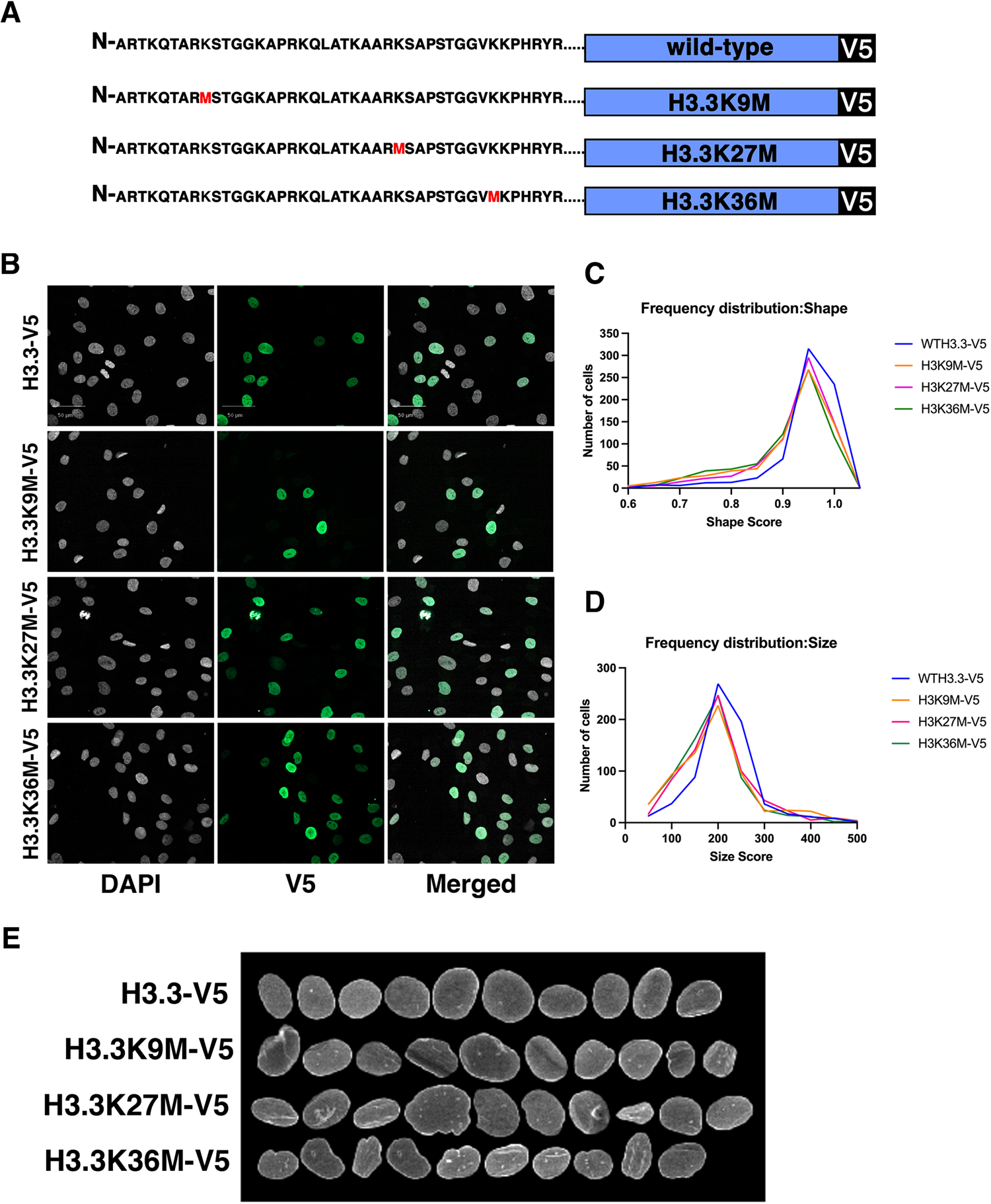
Expression of histone H3.3 mutants affect nuclear shape. (A) WT and mutant histone expression constructs generated for this study. (B) Stable expression of indicated H3.3 mutants in fibroblast cells. Gray: DAPI to detect DNA, Green: V5-tagged histone variant. Scale bar = 50 μm. (C) Frequency distribution of nuclear shape between differing histone H3.3 expression mutants. A p value of <.0001 was obtained for all samples using a two sample Kolmogorow-Smirnov test.

## DISCUSSION

Here we have identified novel determinants of nuclear size and shape by utilizing an imaging based functional genomics screen. Our findings highlight a prominent role of chromatin factors and epigenetic modifiers in the maintenance of nuclear morphology. In support of such a role, using *in vitro* binding assays, we find a direct interaction between lamin A and the modified tail of histone H3 and expression of disease-relevant histone H3.3 mutants altered normal nuclear morphology in human fibroblast cells.

Several cellular factors have been implicated in regulation of nuclear size and shape, including nucleocytoplasmic transport factors and components of the nuclear envelope and nuclear pore complex (Levy and Heald, 2012). In line with these earlier findings, we identified multiple components of nuclear pore complexes such as *NUP205, NUP62, NUPL1,* and *NUP85* as well as a number of nuclear membrane proteins which validates our screening method. Our results are also in line with previous screening studies using the elliptic Fourier coefficient (EFC) as a distinct parameter to quantitatively identify misshapen nuclei in MCF-10A breast epithelial cells which targeted 608 epigenetic gene products and found 33 determinants of nuclear shape, including a number of epigenetic factors (Tamashunas *et al*., 2020). Interestingly, knockdown of some genes encoding core histones such as *HIST1H3B*, *HIST1H4B*, and *HIST1H2BA* also resulted in nuclear morphology defects (Tamashunas *et al*., 2020). Our analysis extends those studies by identifying several epigenetic factors, particularly histone modifiers and readers as determinants of nuclear morphology. Furthermore, our experimental design assessed nuclear size in addition to nuclear shape in multiple cell types. Remarkably, we find distinct sets of size and shape determinants in individual cells lines. Lack of overlap between nuclear size and shape hits even amongst the same cell type underscores the complexity of nuclear morphology regulation and the need for large-scale screens using multiple measurement parameters in parallel to identify regulators of nuclear morphology.

Comparing the size and shape determinants in immortalized human fibroblasts and breast epithelial cells showed remarkably little overlap in determinants of nuclear morphology. The cell-type differences may be due to a number of reasons. One possibility is that different cell types use distinct networks and pathways to regulate nuclear morphology. Given that nuclear morphology does not seem to be controlled by a single dedicated pathway, but rather appears to be the result of multiple mechanisms, this scenario seems unlikely. Alternatively, it is possible that there are innate cellular features amongst cell lines that affect nuclear morphology. For example, differences in the rate of cell division, the amount of cell adhesion, nuclear import/export rates or differing amounts of chromatin or lamin stability may affect nuclear morphology. Previously we found that knockdown of the nuclear pore component ELYS resulted in a nucleus phenotype in breast epithelial cells, and that comparison of nuclear size in ELYS knockdown cells amongst four different cell types found varying degrees of nuclear size reduction (Jevtic *et al*., 2019). These data suggest ELYS functions in regulating nuclear size, although to varying degrees, among differing cell types. In the light of our finding of a prominent role of chromatin and epigenetic factors in determining nuclear morphology, an attractive possibility is that the differences in cellular factors that contribute to nuclear morphology in different cell types reflect cell-type specific epigenetic landscapes in which chromatin modulates nuclear morphology. More systematic analysis of a more diverse set of cell types in future studies should begin to address this question.

Lamins have been widely implicated in maintenance of nuclear morphology (Lammerding *et al*., 2004; Matias et al., 2022). Reassuringly, and as expected, *LMNA* was a hit in our screens. However, many effectors of size and shape identified here exerted their effect without affecting lamin A/C levels. Instead, the nuclear shape screen identified a group of chromatin modifiers. This is in line with previous work pointing to a combined contribution of lamins and chromatin, and their interplay, to nuclear morphology and biophysical properties (Stephens *et al*., 2017). A prominent role of chromatin in nuclear morphology is suggested by the observation that nuclear blebbing can be promoted or inhibited by treating cells with drugs that increase euchromatin or heterochromatin, respectively (Stephens *et al*., 2018). In addition alterations to euchromatin and heterochromatin rescue nuclear morphology defects in disease model cells (Stephens *et al*., 2018). Furthermore, the lysine acetyltransferase HAT1 which acts on newly incorporated histone H4 increased nuclear size, nuclear blebbing and micronuclei and loss of HAT1 acetylation disrupts chromatin regions associated with the nuclear lamina (Popova et al., 2021). Along the same lines, acetylation of lamin A via the acetyltransferase MOF leads to changes in nuclear morphology and epigenetic alterations (Karoutas *et al*., 2019). These observations, combined with our findings, highlight a prominent role of histone modifications in regulation of nuclear size and shape. Our finding of a direct effect of histone modification mutants of H3.3 upon expression *in vivo* support this scenario.

Although it is well established that chromatin is closely juxtaposed with the nuclear lamina and genome regions which associate with the lamina can be mapped as lamin-associated domains (LADs), the precise nature of chromatin-lamin interactions is largely unknown. We find that lamin A, but not lamin C, directly interacts with histone H3. This finding adds to the prior identification of a C-terminal region, present on both lamin A and lamin C, that can bind to DNA, and linker DNA assembled onto nucleosomes (Stierle, 2003). Furthermore, progerin mutations reduced lamin-DNA interactions (Bruston, 2010). Our studies identify a novel lamin A-histone H3 interaction independent of DNA binding. We map two distinct regions located within the C-terminal unstructured tail and within the globular domain which are required for lamin A-H3 interactions and we suggest that these interactions occur in the context of lamin A monomers. The binding of lamin A to histones seems to be facilitated by histone modifications, because we find enhanced binding of lamin A to dual (Rme2s/Kme1-me2) modifications which are associated with transcriptionally repressive marks on heterochromatin (Di Lorenzo and Bedford, 2011; Zhang and Reinberg, 2001). These findings point to a mechanism by which chromatin-lamin A interactions via modified histone H3 tails contribute significantly to nuclear morphology. The precise nature of this interaction and what the downstream effect of this lamin A-histone H3 interaction is will require further studies.

Changes to nuclear shape and size have been documented in many types of cancer and nuclear morphology changes are often correlative with poor prognosis (Pienta and Coffey, 1991; Wolberg et al., 1999; Zink *et al*., 2004). In line with a role of disease-associated histone epigenetic modifiers in contributing to nuclear dysmorphia and disease, we find that expression of K9 and K27 methylation mutants of histone H3.3, in which K27 mutants are also oncogenic, shows changes to nuclear morphology. This finding is relevant since histone H3.3K27M mutations were identified in a subset of pediatric patients with glioblastoma (Khuong-Quang et al., 2012; Schwartzentruber et al., 2012) and H3.3K36M mutations were documented in chondrosarcomas (Behjati et al., 2013), which are also characterized by extensive nuclear aberrations. These findings are in line with our observation of direct physical interaction of lamins with histones.

Taken together, the use of imaging-based screening reported here significantly expands the list of cellular factors that contribute to nuclear morphology. Our findings of an enrichment of chromatin factors and the fact that the vast majority of nuclear size and shape effectors exerts their function without alteration of lamin protein levels highlights the important role chromatin plays in determining nuclear shape and size.

## Acknowledgements

We would like to thank Dr. Leonard Kubben, Ph.D. (IMB, Mainz), Dr. Sigal Shachar, Ph.D., and Dr. Akanksha Singh, Ph.D. (Active Motif) for advice, protocols, and troubleshooting. Work in the Misteli lab and at HiTIF was supported by the Intramural Research Program of the NIH, NCI, Center for Cancer Research via 1-ZIA-BC010309-23 and 1-ZIC-BC011567-08, respectively. Work in the Levy lab was supported by the National Institutes of Health/National Institute of General Medical Sciences (R35GM134885 and P20GM103432) and the USDA National Institute of Food and Agriculture (Hatch project #1012152).

---

**SUPPLEMENTAL FIGURE 1:**
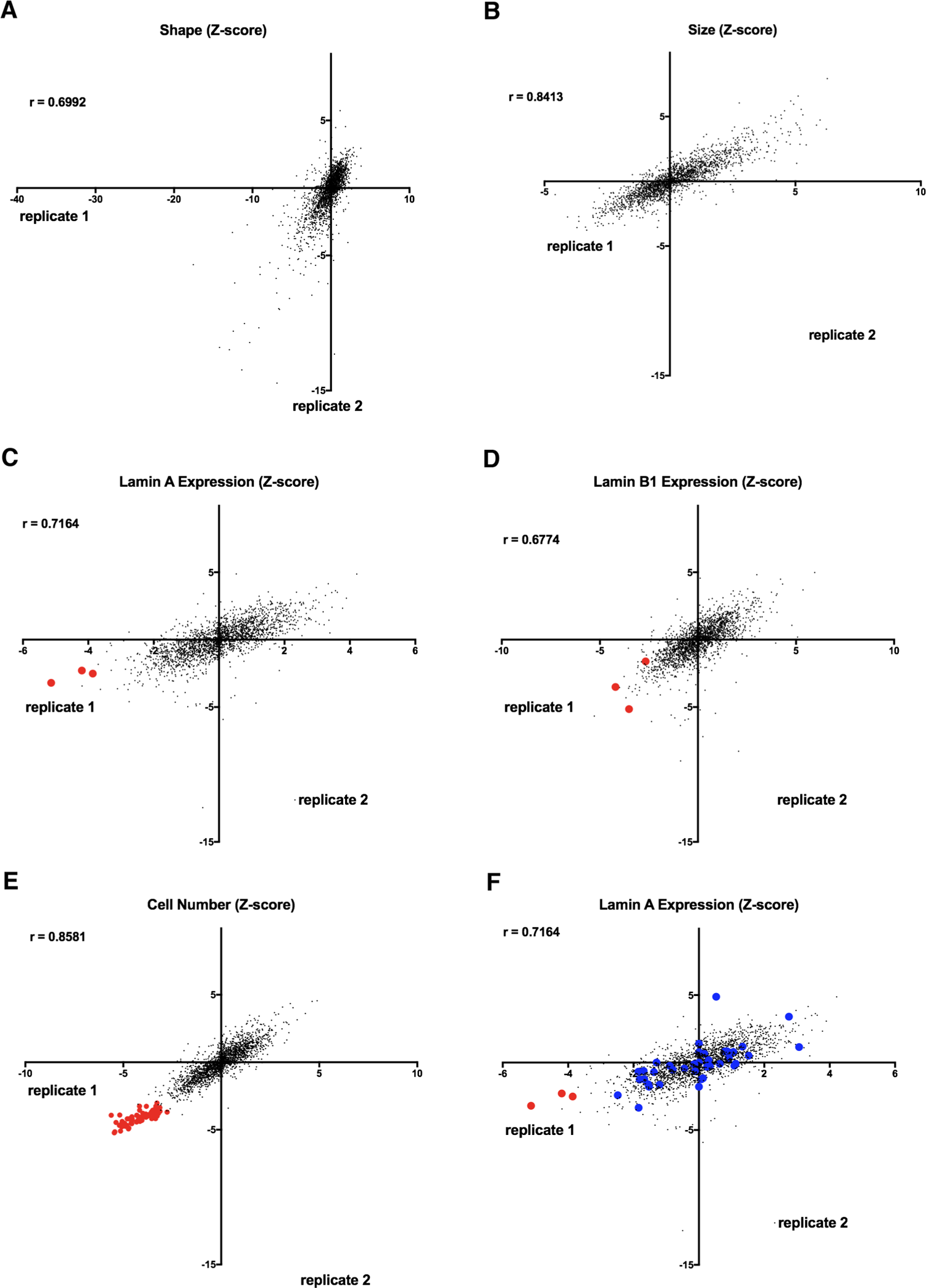
Reproducibility of nuclear morphology screen using human fibroblast cells. (A) Nuclear shape z-scores of the two replicates (Replicate 1 and Replicate 2) of the nuclear shape screen. (B) Nuclear size z-scores of the two replicates (Replicate 1 and Replicate 2) of the nuclear shape screen. (C) Lamin A/C expression z-scores of the two replicates (Replicate 1 and Replicate 2) of the nuclear shape screen. Red dots indicate siRNAs targeting *LMNA* gene products. (D) Lamin B1 expression z-scores of the two replicates (Replicate 1 and Replicate 2) of the nuclear shape screen. Red dots indicate siRNAs targeting *LMNB1* gene products. (E) Correlation plot shows cell number z-scores of the two replicates (Replicate 1 and Replicate 2) of the nuclear shape screen. Red dots indicate siRNAs designed to cause cell death as an indicator of transfection efficiency. (F) Correlation plot shows lamin A/C expression z-scores of the two replicates (Replicate 1 and Replicate 2) of the nuclear shape screen. Data points labeled in red indicate siRNAS targeting *LMNA* gene products while shape hits are highlighted in blue.

**SUPPLEMENTAL FIGURE 2:**
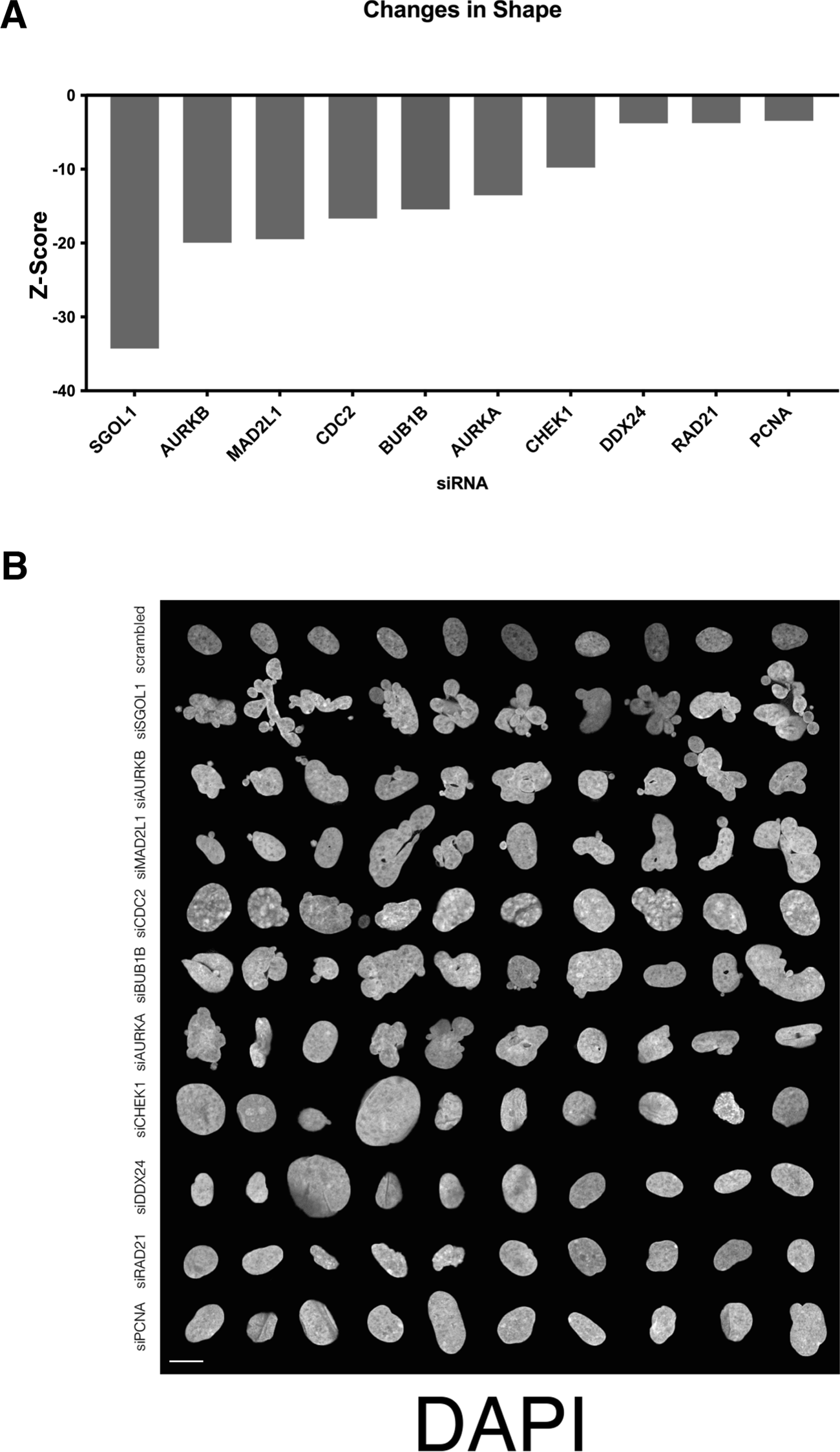
Nuclear shape hits that affected cell number. (A) Z-scores of the top nuclear shape determinants which had low cell number. (B) Montage of nuclear shape determinants with low cell number revealed misshapen nuclei and mitotic defects. Signal represents DAPI staining. Scale bar = 10 μm.

**SUPPLEMENTAL FIGURE 3:**
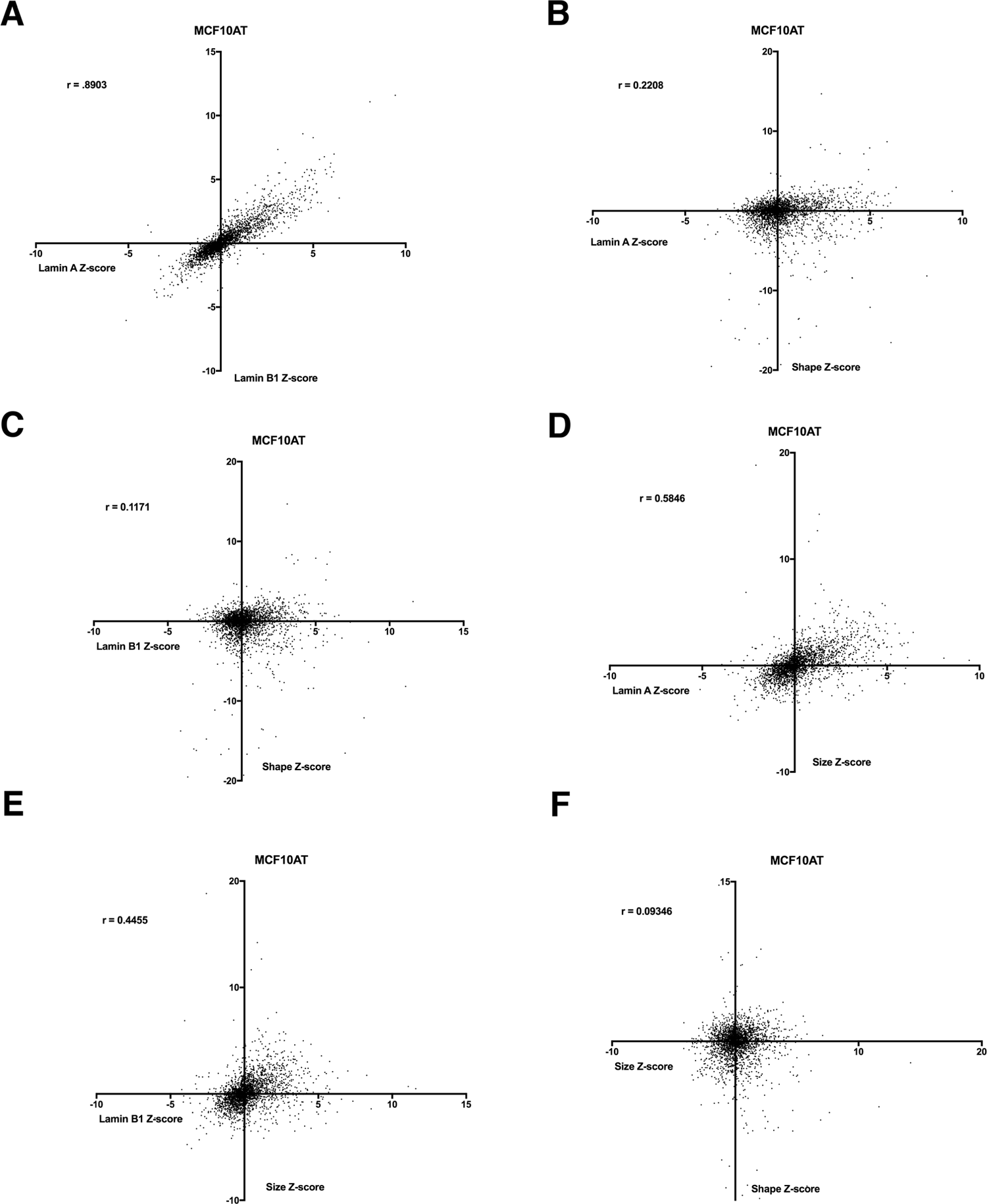
Correlation plots of nuclear morphology data using MCF10AT human breast epithelial cells. (A) Lamin A/C expression z-scores compared to Lamin B1 z-scores in MCF10AT cells. (B) Lamin A/C expression z-scores compared to nuclear roundness z-scores in MCF10AT cells. (C) Correlation plot of Lamin B1 expression z-scores relative to nuclear shape z-scores in MCF10AT cells. (D) Lamin A/C expression z-scores compared to nuclear size z-scores in MCF10AT cells. (E) Lamin B1 expression z-scores compared to nuclear size z-scores in MCF10AT cells. (F) Correlation plot of nuclear shape z-scores compared to nuclear size z-scores in MCF10AT cells.

**SUPPLEMENTAL FIGURE 4:**
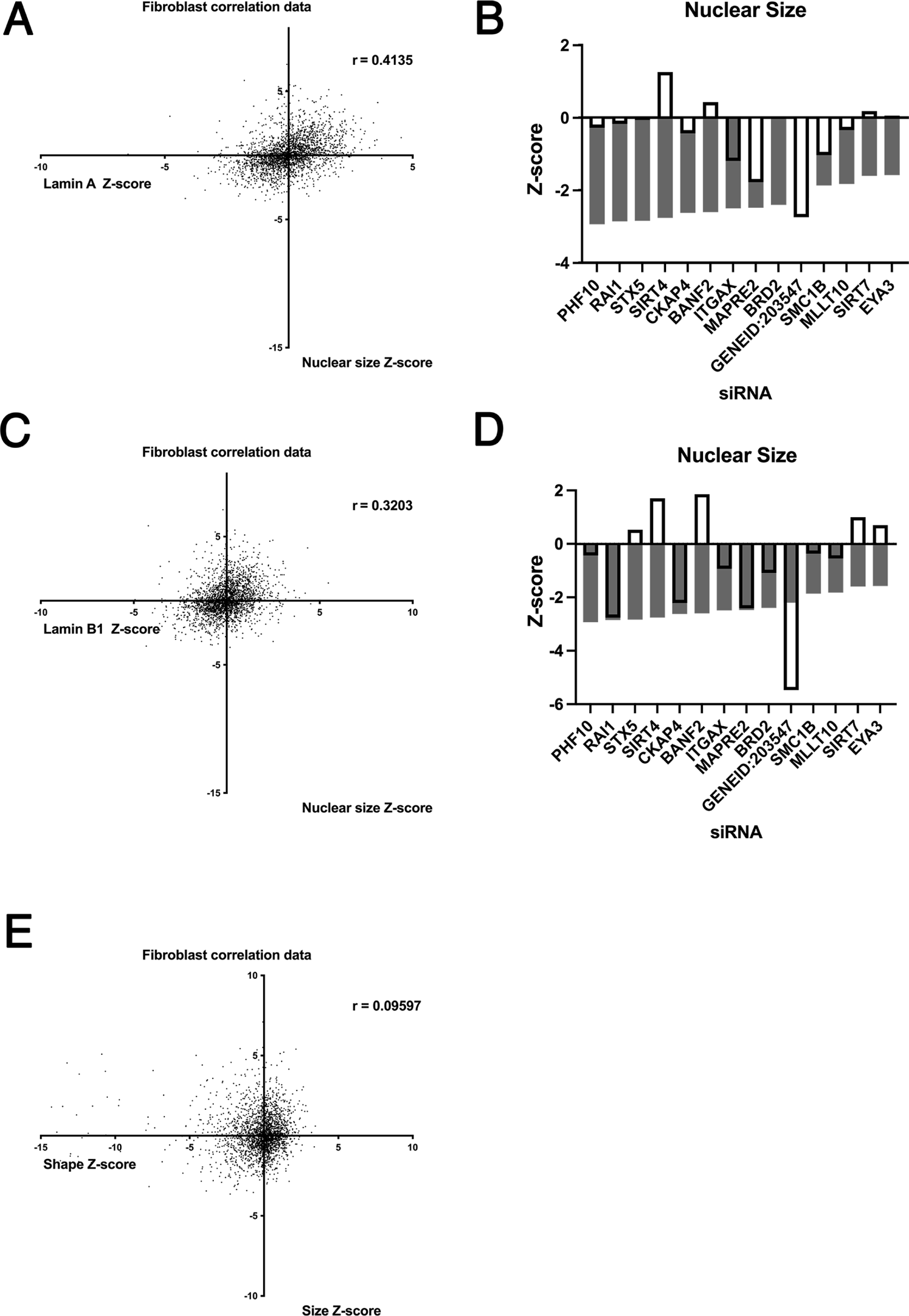
Nuclear morphology data using human fibroblast cells reveal lack of correlation between lamin expression and nuclear morphology features. (A) Lamin A/C expression z-scores compared to nuclear size z-scores in human fibroblast cells. (B) Nuclear size z-scores (gray bars) compared to lamin A/C expression (bars with black outline) reveals little correlation between lamin A/C expression and decreased nuclear size. (C) Correlation plot comparing lamin B1 expression z-scores to nuclear size z-scores in human fibroblast cells. (D) Nuclear size z-scores (gray bars) compared to lamin B1 expression z-scores (bars with black outline) reveals little correlation between lamin B1 expression and decreased nuclear size. (E) Nuclear shape z-scores compared to nuclear size z-scores in human fibroblast cells.

**SUPPLEMENTAL FIGURE 5:**
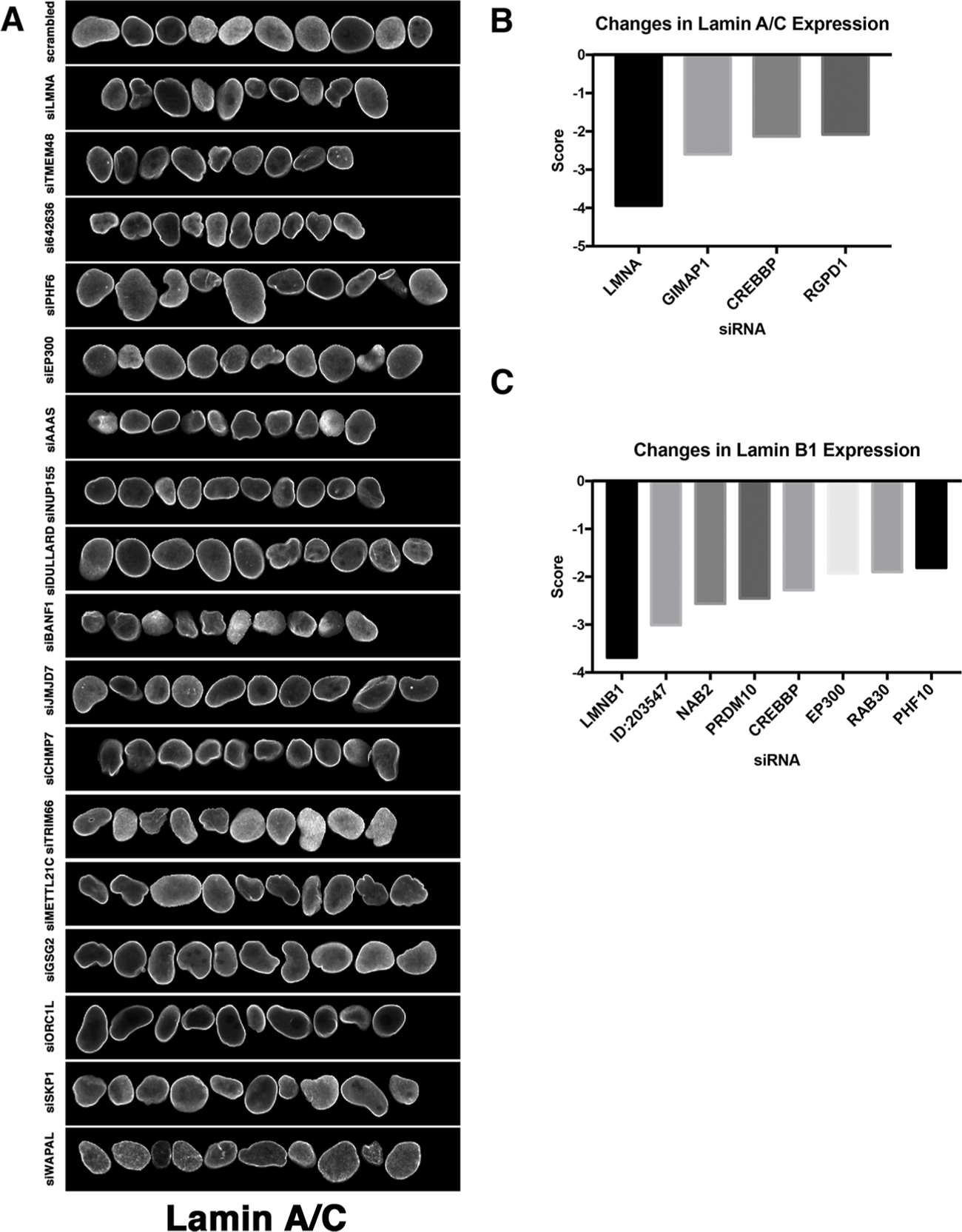
Identification of nuclear shape determinants in MCF10AT cells. (A) Representation of normal nuclei and nuclear shape hits identified by high-throughput screening in MCF10AT cells. Nuclear shape abnormalities are visualized by lamin A/C antibody staining. Scale bar = 10μm. (B) Nuclear intensity of lamin A/C was assessed by calculating Z-scores of changes in lamin A/C expression on a per well basis. Lamin A/C hits were identified by z-scores of −1.5 or less. (C) Nuclear intensity of lamin B1 was assessed by calculating z-scores of changes in lamin B1 expression of −1.5 or less per well.

**SUPPLEMENTAL FIGURE 6:**
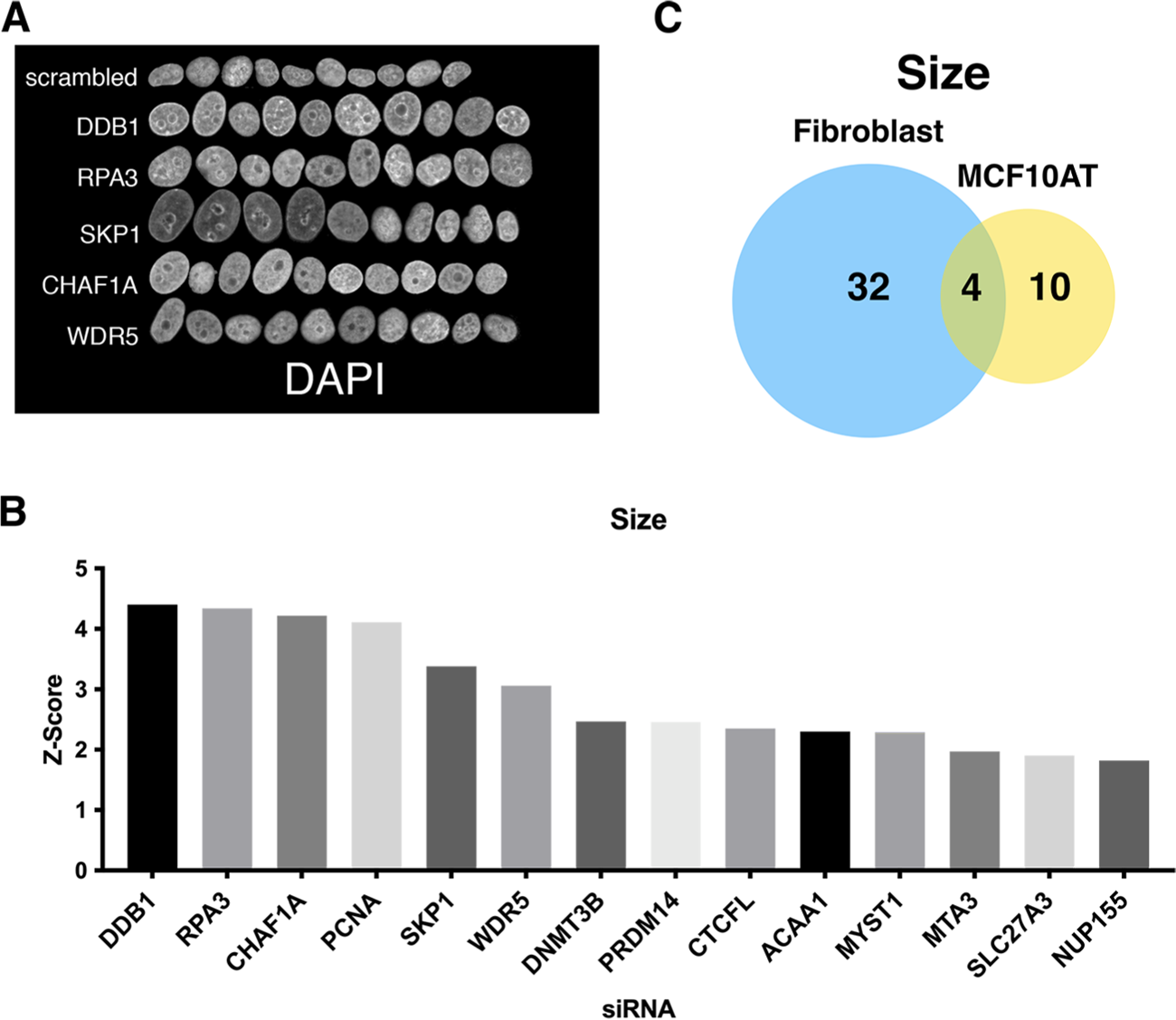
Identification of nuclear size determinants in MCF10AT cells. (A) Representation of normal nuclei and hits with enlarged nuclei identified by high-throughput screening in MCF10AT cells. Nuclear size abnormalities are visualized by DAPI staining. Scale bar = 10μm. (B) Nuclear size hits were calculated by scoring nuclear area. z-scores were generated to compare hits across the screen. Hits were identified as having a z-score of −1.5 or less. At least 250 nuclei were analyzed per sample. (C) Little overlap of hits for nuclear size changes in immortalized human fibroblast cells compared to nuclear size hits for the breast epithelial cell line MCF10AT.

**SUPPLEMENTAL FIGURE 7:**
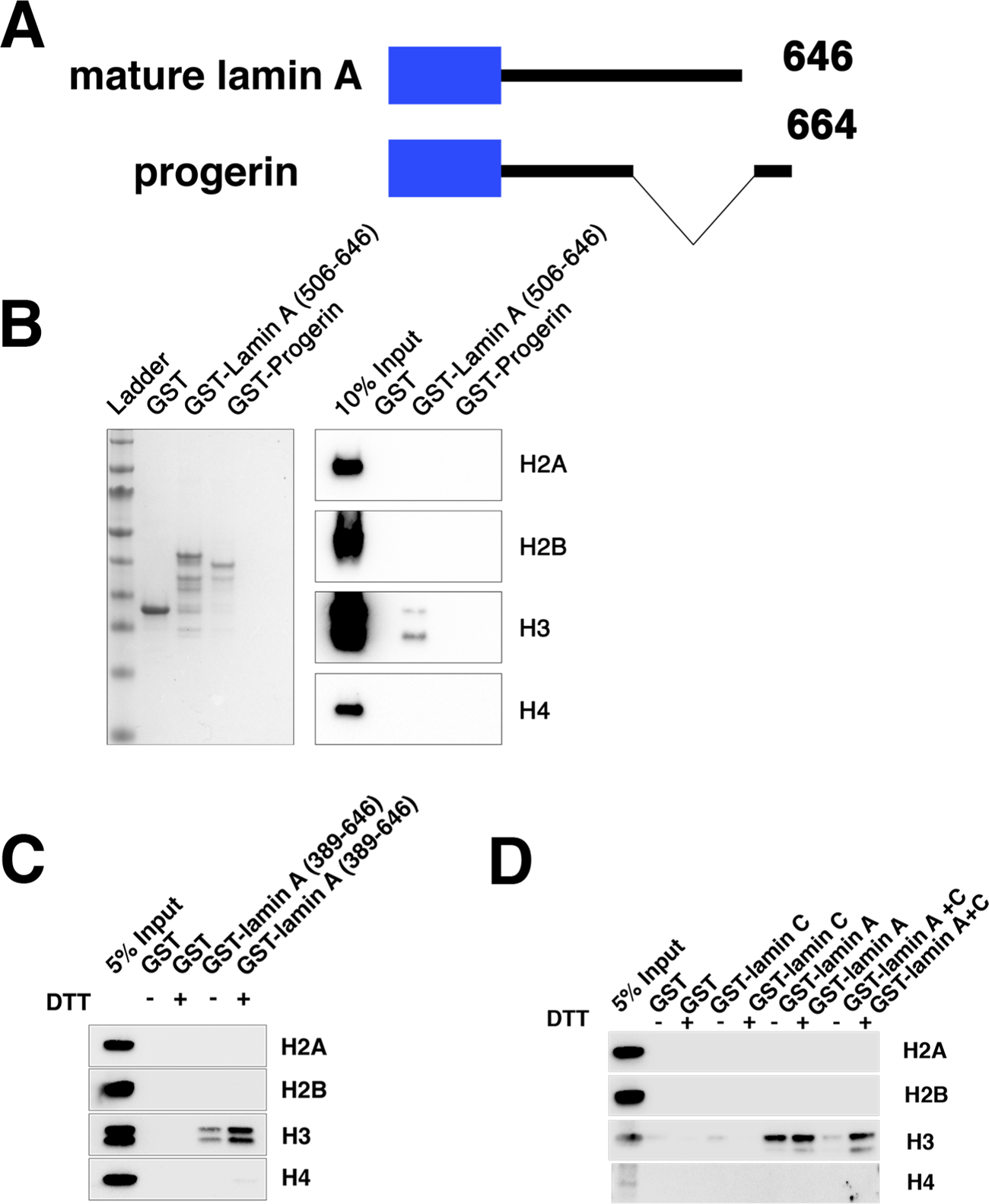
Lamin C inhibits lamin A-H3 interactions. (A) A diagram of mature lamin A and progerin constructs used in the binding assays. (B) A GST pull-down assay with wild-type GST-lamin A and a mutated lamin A (GST-Progerin) encoding the disease causing lamin A construct showed that the recombinant progerin peptide could not bind to histone H3. (C) A GST pull-down assay identified that the addition of a reducing agent (DTT) which inhibits lamin A dimerization promotes lamin A-histone H3 interactions. (D) A GST pull-down assay mixing lamin C and lamin A constructs in the presence of DTT identified lamin A-H3 interactions are inhibited by the addition of lamin C and this can be alleviated by adding DTT.

**SUPPLEMENTAL FIGURE 8:**
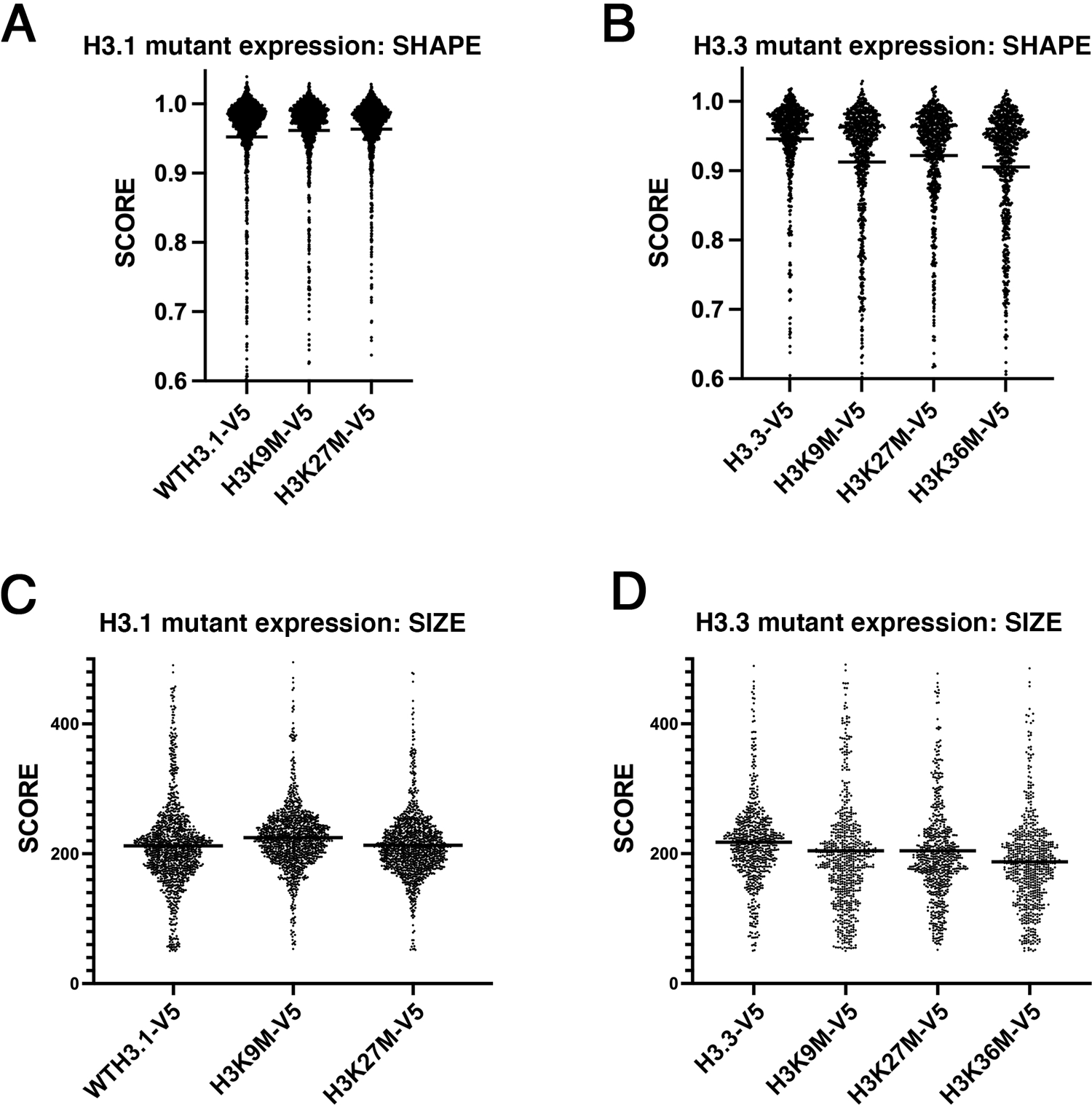
Histone H3.1 mutants display a lesser nuclear morphology phenotype compared to histone H3.3 mutants. (A) Cells expressing wild-type histone H3.1, H3.1K9M or H3.1K27M mutants showed little change in nuclear shape scores. The mean is indicated by the horizontal line. (B) Cells expressing histone H3.3 constructs reveal histone H3.3K9, H3.3K27M, or H3.3K36M mutants showed reduced nuclear shape scores compared to wild-type H3.3 expression. The mean is indicated by the horizontal line. (C) Cells expressing wild-type histone H3.1, H3.1K9M or H3.1K27M mutants showed some change in nuclear size scores. The mean is indicated by the horizontal line. (D) Cells expressing histone H3.3 constructs reveal histone H3.3K9, H3.3K27M, or H3.3K36M mutants showed reduced nuclear size scores compared to wild-type H3.3 expression. The mean is indicated by the horizontal line.

**TABLE S1:**
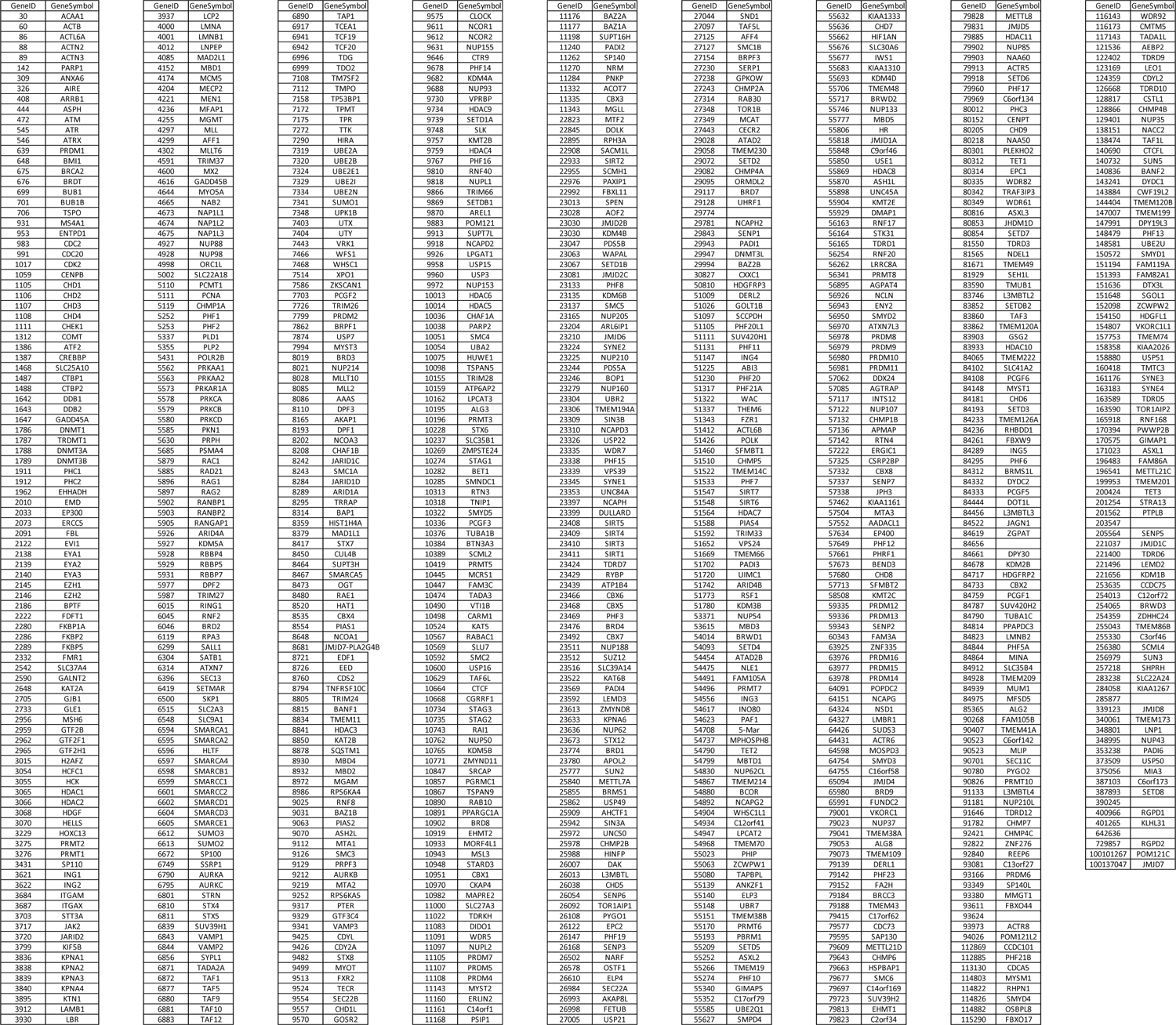
TARGET GENE LIST

**TABLE S2:** Plasmids used in this study

| In vitro protein expression plasmids |  |  |
| --- | --- | --- |
| Plasmid Name: | Content: | Cloning Sites: |
| pACS34 | GST (PGEX4T1) |  |
| pACS45 | GST-lamin A (389-646) | BAMHI/ECORI |
| pACS57 | GST-lamin C (389-572) | BAMHI/XHOI |
| pACS58 | GST-lamin A (565-646) | BAMHI/ECORI |
| pACS69 | GST-lamin A (389-566) | BAMHI/ECORI |
| pACS77 | GST-lamin A (389-638) | BAMHI/ECORI |
| pACS75 | GST-lamin A (389-626) | BAMHI/ECORI |
| pACS73 | GST-lamin A (389-609) | BAMHI/ECORI |
| pACS71 | GST-lamin A (389-597) | BAMHI/ECORI |
| pACS37 | GST-lamin A (506-646) | BAMHI/XHOI |
| pACS38 | GST-Progerin (506- (506-664 delta 50) | BAMHI/XHOI |
| pACS122 | GST-lamin A (451-646) | BAMHI/ECORI |
| pACS124 | GST-lamin A (551-646) | BAMHI/ECORI |

**Table S3:** Antibodies used in this study:

| Target Antigen: | Company: | Catalog # | Concentration: |
| --- | --- | --- | --- |
| Histone H2A | Abcam | AB18255 | 1:250 |
| Histone H2B | Abcam | AB1790 | 1:1000 |
| Histone H3 | Abcam | AB1791 | 1:5000 |
| Histone H4 | Abcam | AB16254 | 1:1000 |
| Lamin A/C | SantaCruz | SC376248 | 1:1000 |
| Lamin B1 | SantaCruz | SC-6217 | 1:500 |
| Lamin B1 | Abcam | AB16048 | 1:1000 |

**TABLE S4A:**
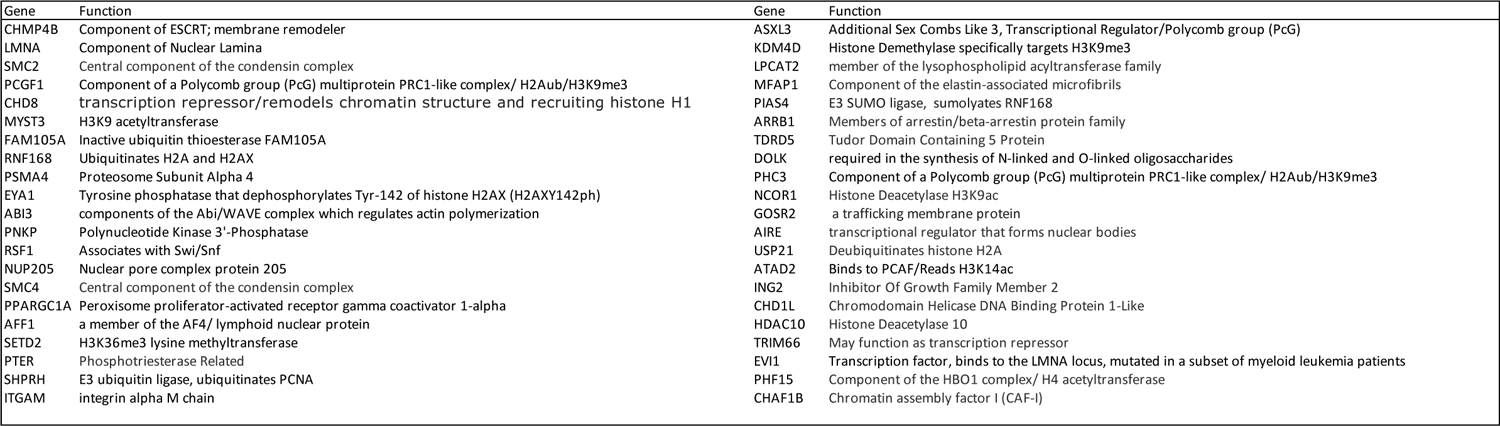
Nuclear Shape Hits FIBROBLAST

**TABLE S4B:**
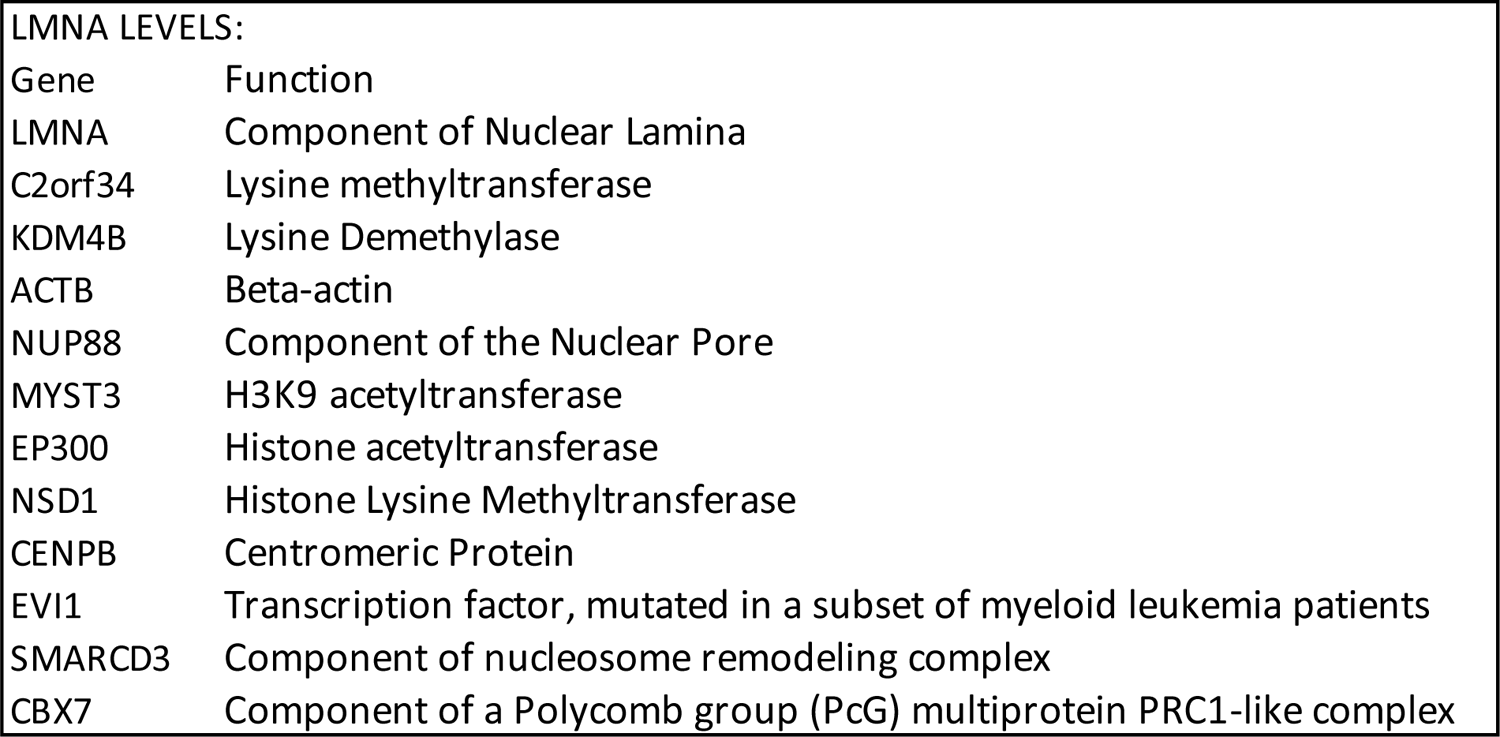
LMNA Expression Hits-FIBROBLAST

**TABLE S4C:**
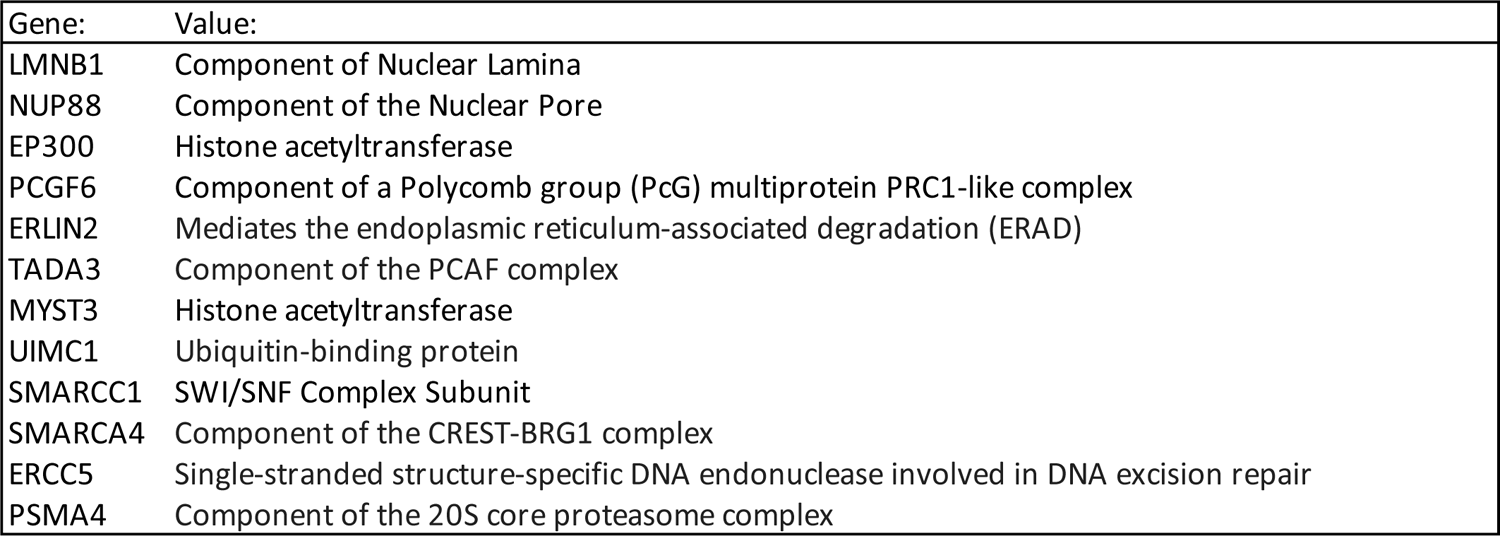
LMNB1 Expression Hits - FIBROBLAST

| Gene: | Value: |
| --- | --- |
| LMNB1 | Component of Nuclear Lamina |
| NUP88 | Component of the Nuclear Pore |
| EP300 | Histone acetyltransferase |
| PCGF6 | Component of a Polycomb group (PcG) multiprotein PRC1-like complex |
| ERLIN2 | Mediates the endoplasmic reticulum-associated degradation (ERAD) |
| TADA3 | Component of the PCAF complex |
| MYST3 | Histone acetyltransferase |
| UIMC1 | Ubiquitin-binding protein |
| SMARCC1 | SWI/SNF Complex Subunit |
| SMARCA4 | Component of the CREST-BRG1 complex |
| ERCC5 | Single-stranded structure-specific DNA endonuclease involved in DNA excision repair |
| PSMA4 | Component of the 20S core proteasome complex |

**TABLE S5:**
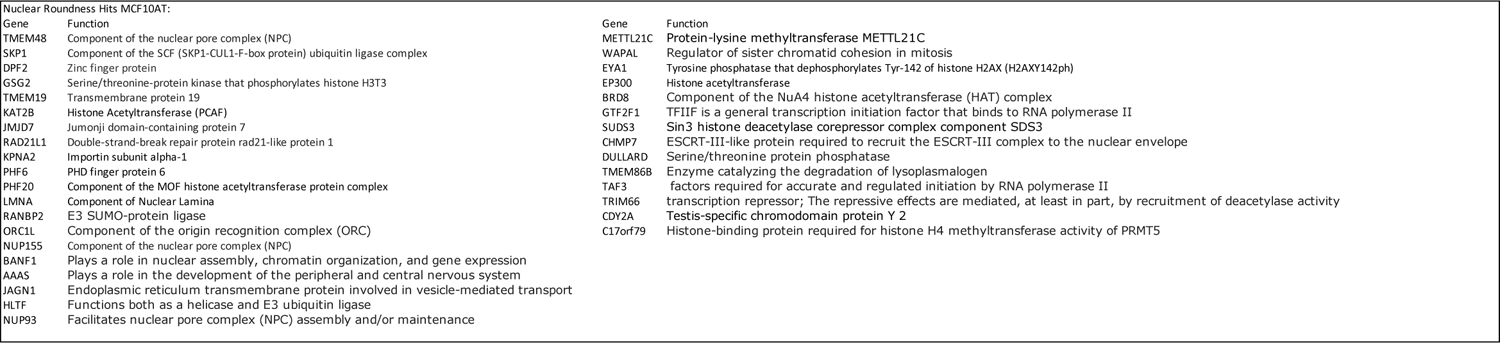
MCF10AT Shape Hits

**TABLE S6:** Top Lamin A interacting peptides

| Rank | Name | Mod 1 | Mod 2 | Mod 3 | Mod 4 | Intensity |
| --- | --- | --- | --- | --- | --- | --- |
| 1 | H3 1-19 | R2me2a | T3P | K4me3 |  | 0.85037936 |
| 2 | H3 1-19 | R8me2s | K9me2 |  |  | 0.61669194 |
| 3 | H3 16-35 | R26me2s | K27me2 |  |  | 0.6030349 |
| 4 | H4 11-30 | R19me2s | K20me1 |  |  | 0.59393018 |
| 5 | H3 1-19 | R2me2s | K4me3 |  |  | 0.54840667 |
| 6 | H3 1-19 | R2me2a | K4me1 |  |  | 0.51411228 |
| 7 | H4 11-30 | K12ac | K16ac |  |  | 0.49590287 |
| 8 | H3 1-19 | R2me2a | K4me1 | R8me2s | K9me2 | 0.4488619 |
| 9 | H3 1-19 | R2me2a | K4me1 | K9ac |  | 0.44339908 |
| 10 | H3 1-19 | K9me2 |  |  |  | 0.43338389 |
| 11 | H4 11-30 | K16ac | R17me2s |  |  | 0.4151745 |
| 12 | H3 16-35 | K27ac |  |  |  | 0.41305006 |
| 13 | H3 1-19 | R2me2a | K4ac | R8me2s | K9me2 | 0.37481031 |
| 14 | H3 1-19 | K4me1 |  |  |  | 0.36631259 |
| 15 | H3 16-35 | R26me2a | K27ac |  |  | 0.35872533 |
| 16 |  | Background 1 |  |  |  | 0.35386948 |
| 17 | H3 1-19 | K4ac |  |  |  | 0.34962062 |
| 18 | H3 1-19 | R2me2s | K4me1 |  |  | 0.34901364 |
| 19 | H3 1-19 | R2me2s | K4me2 | R8me2s | K9me2 | 0.34385431 |
| 20 | H2b 1-19 | K5ac | K12ac | K15ac |  | 0.34355082 |
| 21 | H3 1-19 | K4me2 | R8me2s | K9me3 |  | 0.32503792 |
| 22 | H3 1-19 | T3P |  |  |  | 0.29681334 |
| 23 | H3 1-19 | R2me2s | K4me1 | R8me2s | K9me2 | 0.2855842 |
| 24 | H3 1-19 | R2me2a | K4me2 | R8me2s | K9me2 | 0.28406675 |
| 25 | H3 1-19 | R2me2a | T3P | K4me2 |  | 0.28285279 |

